# Species-specific metabolic networks shape evolutionary routes to functional rescue

**DOI:** 10.64898/2026.09.15.751699

**Authors:** Shanni Hornstein, Swati Sirotiya, Raul Mireles, Orr Giladi, Alexander Geller, Carolina Cano-Prieto, Daniela Rago, Linda Ahonen, Tamar Cohen Shapir, Amichai Baichman-Kass, Pablo Cruz-Morales, Asaf Levy, Lianet Noda-García

**Author notes:** These authors contributed equally to this work (Shanni Hornstein, Swati Sirotiya).

## Abstract

Metabolic networks are highly interconnected. Still, it remains unclear whether organism-and environment-specific factors shape their capacity to evolve in response to metabolic stress or whether it follows general principles. Here, we studied metabolic evolvability in *Bacillus subtilis* using auxotrophic mutants lacking central biosynthetic enzymes. Across two genetic backgrounds with contrasting biofilm-forming capacities and under direct or gradual selection through nutrient gradients, *B. subtilis* bypassed 9 of 17 essential functions. Rescue was more frequent in the biofilm-proficient background and under gradual selection, which was also associated with more mutations in coding regions, particularly nonsynonymous. Adaptation proceeded through loss-of-function mutations that relieved regulatory or enzymatic constraints and redirected metabolic flux. Comparison with *Escherichia coli* revealed substantial differences in bypassability and genetic routes that persisted under matched conditions, although two cross-species solutions converged at the pathway level. Our data show species-specific metabolic networks shape available rescue routes, while ecological context influences their evolutionary accessibility during adaptation.

## Introduction

Metabolic networks exhibit robustness and plasticity, allowing organisms to tolerate genetic and environmental perturbations ^1–4^. This capacity emerges from network connectivity, functional redundancy, and underground metabolism (UM), a reservoir of latent catalytic activities and metabolic routes that normally carry little or no physiologically relevant flux ^5,6^. Under severe genetic perturbations, these activities can restore metabolic flux and support survival ^7–9^. Understanding which metabolic functions can be bypassed and through which evolutionary routes is therefore central to understanding metabolic robustness and evolvability.

Much of our understanding of metabolic rescue after gene loss comes from systematic studies in *Escherichia coli*. Gene-deletion collections, multicopy suppression screens, and adaptive laboratory evolution (ALE) have revealed compensation mechanisms for missing metabolic functions ^10,11^. These include recruiting and optimizing promiscuous enzyme activities, activating alternative pathways, and redirecting flux through regulatory changes. They arise through coding or regulatory mutations and copy-number amplifications ^7,9,12,13^. Experimental evolution further suggests that responses to metabolic gene loss are repetitive and thus predictable ^13^. Notably, three independent studies of disrupted IlvA-dependent isoleucine biosynthesis converged on sulfur amino-acid metabolism, including genes associated with methionine and cysteine biosynthesis ^13–15^.

Collectively, this body of work has established *E. coli* as the primary model for understanding how metabolic networks adapt to the loss of individual functions.

Whether these principles represent general properties of bacterial metabolism, however, remains unclear. Computational comparisons indicate that perturbing a conserved metabolic reaction can have species-specific consequences depending on the network context ^16,17^. Likewise, the accessible underground network should depend on each species’ enzyme repertoire, secondary activities, and regulatory architecture, shaped by lineage-specific evolutionary history ^18,19^. Thus, even conserved metabolic functions may therefore differ among species in their capacity for functional rescue and the genetic routes by which alternative solutions become accessible. Environmental context may add variation. Most systematic ALE studies of metabolic rescue have used homogeneous, well-mixed cultures ^7,13,14^, whereas microorganisms frequently inhabit spatially structured environments. Biofilms and nutrient gradients create heterogeneous conditions that can sustain phenotypic and genetic diversity and alter evolutionary trajectories ^20,21^. Distinguishing species-specific metabolic constraints from environmental effects is therefore essential for determining how broadly rescue evolutionary routes can be generalized.

*Bacillus subtilis* provides an experimentally tractable, phylogenetically distant counterpoint to *E. coli*. Its characterized metabolism and genome-wide deletion collection make it well suited for systematic analysis of metabolic gene loss ^22,23^, while its planktonic and structured biofilm lifestyles allow evolution to be examined under contrasting ecological conditions ^24^. Here, we investigate *B. subtilis’* capability to overcome 17 essential-gene deletions across two genetic backgrounds differing in biofilm-forming capacity and under homogeneous or nutrient-gradient selection. By combining ALE, whole-genome sequencing, metabolomics, proteomics, and biochemical characterization, we identify rescued functions and their mechanisms. Comparison with published *E. coli* evolution experiments and matched structured-condition experiments reveals substantial differences in the accessibility and genetic basis of functional rescue, alongside specific metabolic solutions shared across species. Our results indicate that species-specific metabolic network properties shape evolutionary routes to functional rescue, while ecological context modulates access to these solutions.

## Results

### 1. Systematic evolution reveals extensive but uneven functional rescue in *B. subtilis*

To explore the capacity of the *B. subtilis* metabolic network to compensate for the loss of essential biosynthetic functions, we selected 17 auxotrophic strains from the *B. subtilis* 168 gene knockout collection (Table S1). Each carried a kanamycin resistance cassette replacing a gene involved in the biosynthesis of an amino acid or other essential metabolite ^22^. Auxotrophy was confirmed in chemically defined minimal medium containing glycerol and glutamate as carbon and nitrogen sources (MSgg). None grew in MSgg after 10 days, whereas all grew in rich medium (LB supplemented with Kanamycin). We reconstructed the same 17 deletions in the *B. subtilis* NCIB3610 background, which, unlike the 168 background, readily forms structured biofilms in MSgg ^25^. Similarly, we confirmed their auxotrophy, generating 34 deletion strains: 17 metabolic functions in two genetic backgrounds.

We subjected these deletion strains to long-term selection for functional rescue (Figure 1A) either directly onto MSgg agar or onto nutrient-gradient plates (Figure S1), providing a gradual transition from permissive to selective conditions. We incubated plates for up to three months. When growth was detected, the population was recovered and genotyped to confirm the original deletion and exclude cross-contamination. Populations were considered successfully rescued only when growth was maintained through at least two consecutive cycles in MSgg.

**Figure 1.**
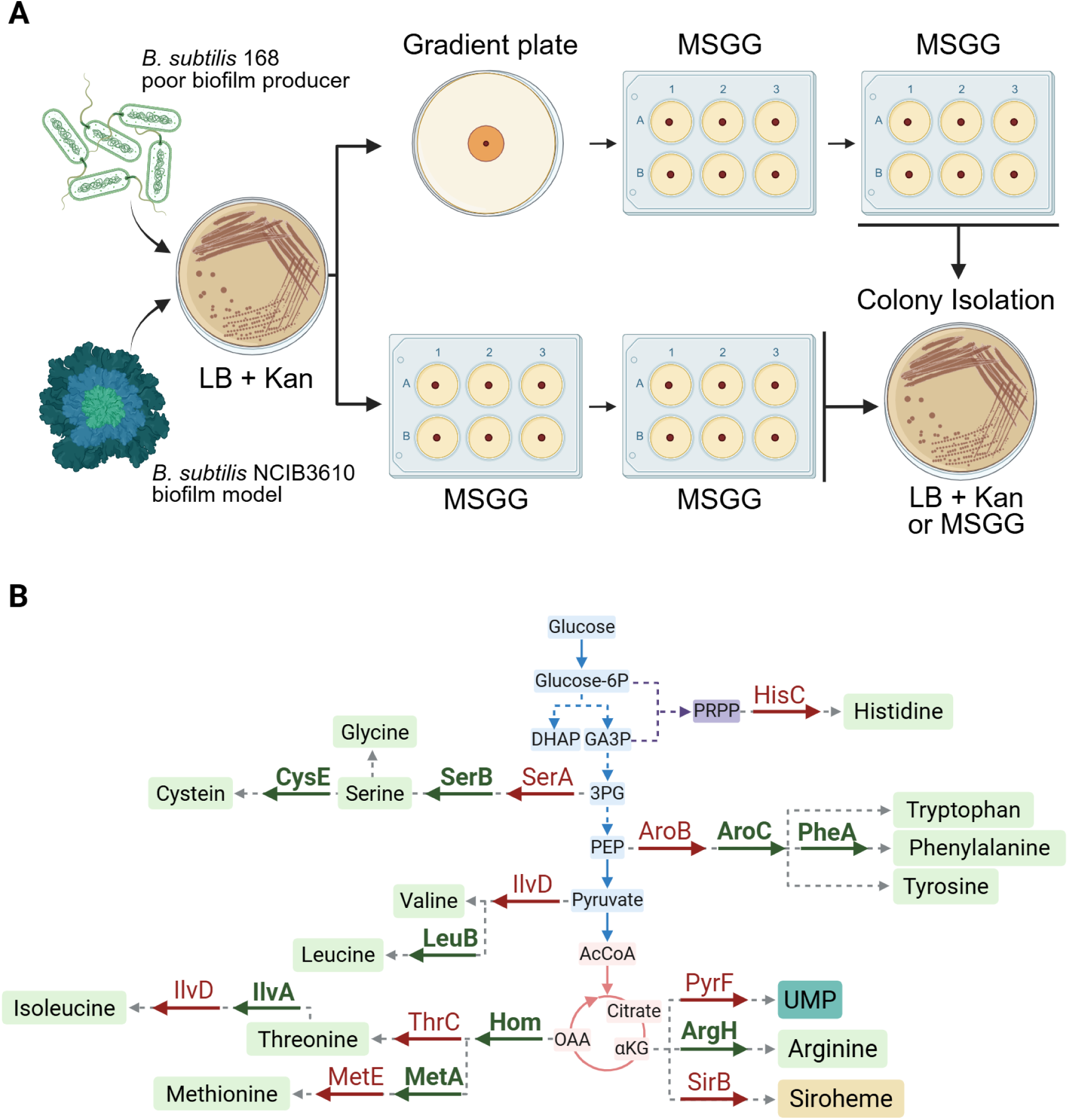
*B. subtilis* can evolutionarily bypass approximately half of the central metabolic functions tested. **(A)** Experimental framework for systematic selection of metabolic functional rescue. **(B)** Evolutionary rescue bypasses 9 of 17 central metabolic functions (*ilvD* is shown twice as it performs the same reaction in two pathways). Enzymes are named by their 4-letter code, as used in Subtiwiki ^32^. Green names and arrows indicate functions that were successfully bypassed. Red indicates functions that were not bypassed. Glycolytic intermediates are shown in blue boxes, tricarboxylic acid cycle intermediates in pink boxes, and pentose phosphate intermediates in purple boxes. Final products, mainly amino acids, are shown in green boxes; UMP and Siroheme, corresponding to pyrimidines and cofactor biosynthesis, are shown in cyan and yellow boxes, respectively. The metabolic map is simplified, and not all connections are shown. Metabolite and enzyme names are provided in Table S1.

For each deletion strain, we performed 2–8 independent evolution experiments, totaling 242 experiments. Of these, 46 populations (19%) representing 16 of the 34 deletion strains, evolved to sustain growth in MSgg and bypassed 9 of the 17 functions (Figure 1B). The functions encoded by *argH, aroC, cysE, ilvA, leuB, pheA*, and *serB* were bypassed in both genetic backgrounds, whereas *hom* and *metA* were bypassed only in NCIB3610. In contrast, *aroB, hisC, ilvD, metE, pyrF, serA, sirB*, and *thrC* were never bypassed (Table S2). Visible growth appeared after 4-93 days (Figure S3; Table S2). Growth time varied widely across functions, and slowly growing populations were subjected to additional MSgg passages to confirm rescue. (Figure S4). Finally, after two or more consecutive MSgg passages, we isolated two colonies from each population.

We next asked whether bypassability could be explained by simple measures of genomic redundancy. Paralogues or enzymes catalyzing similar chemical transformations could provide alternative activities ^26–31^. Thus, we compared the number of paralogues and enzymes with the same first three digits of the Enzyme Commission (EC) classification between bypassed and non-bypassed functions. Neither measure differed significantly (*P* > 0.1; Table S3; Figure S2).

Thus, *B. subtilis* can evolve in the absence of approximately half of the biosynthetic functions examined, but bypassability is unevenly distributed and not predicted by simple measures of genomic redundancy.

### 2. Functional rescue occurs through diverse genetic trajectories

To identify genetic changes underlying functional rescue, we whole-genome sequenced 46 populations, two colonies per population, and 16 corresponding ancestors to ∼200× coverage Breseq ^33^ was run in polymorphism mode for populations (retaining mutations >25% estimated frequency) and in clonal mode for evolved and ancestral colonies (using the default 80% frequency cutoff; consensus mutations are reported as 100%) The *Bacillus subtilis* 168 and NCIB3610 genomes (NC_000964 and NZ_CP020102.1, respectively) served as references and were reannotated using RAST ^34^ to standardize protein-coding sequence names. Mutations present in the ancestors were removed, yielding a final list of plausible evolution-acquired mutations (Table S4). Half the colonies were also analyzed using Snippy ^35^, which matched Breseq for 95% of mutations (Table S4), supporting the validity of the mutation list.

Most populations (34/46, 73.9%) and colonies (78/92, 84.7%) carried one to nine mutations. Two populations and five colonies were hypermutators, accumulating 25–76 mutations (Figure 2A), whereas ten populations and nine colonies had no detectable mutations. These were predominantly colonies bypassing *pheA*, some of which grew within ∼14 days. This is consistent with reports that *pheA* deletion strains can grow without phenylalanine supplementation, suggesting that alternative routes to phenylalanine production may exist ^36^.

**Figure 2.**
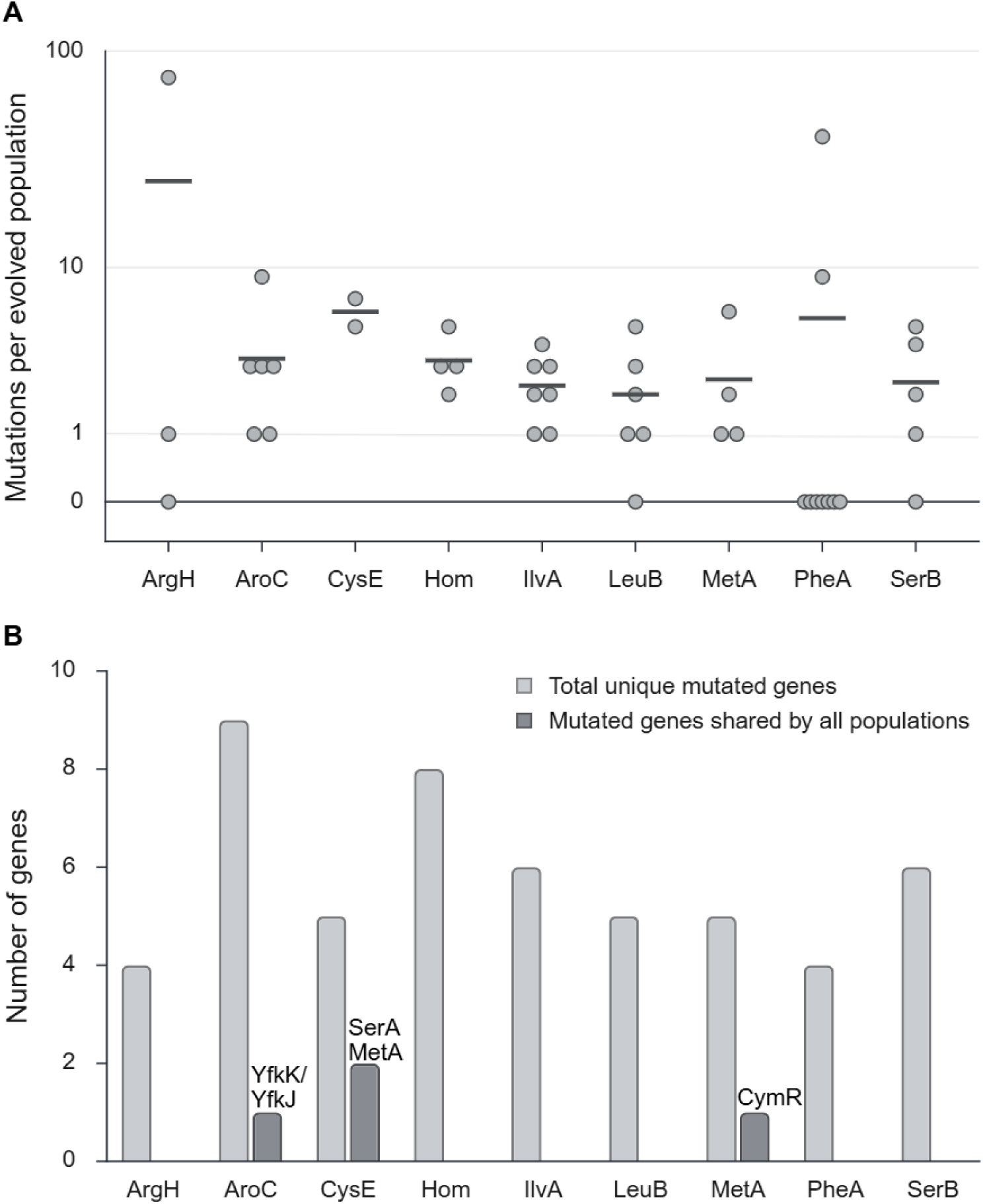
Mutation analysis showed diverse genetic trajectories in functional rescue. **(A)** Mutation count per function. The x-axis shows the missing functions, and the y-axis shows the total number of mutations in each population (log scale). Each dot represents a population missing the indicated function, and the horizontal line represents the mean number of mutations among populations with the same deletion. **(B)** Mutational convergence among independent populations bypassing the same function. The x-axis shows the missing functions, and the y-axis shows the total number of mutated genes across populations missing the same function. Unique mutated genes are in light gray, and genes shared by all populations are in dark gray: AroC: YfkK/YfkJ; CysE: SerA, MetA; MetA: CymR.

Excluding hypermutators, we identified 306 mutations: 75% single-nucleotide polymorphisms, 23% small insertions or deletions, and 2% larger deletions. Copy-number variation affected 11 regions spanning 5–296 kb in 20 colonies (Table S4). Overall, 76% of population mutations were also detected in corresponding colonies, although correspondence varied by function and was particularly low for *argH*.

We next searched for mutational convergence among independent populations bypassing the same function, reasoning, like others ^37,38^, that recurrent mutations in the same gene could indicate a functional role. For *aroC*, *cysE*, and *metA*, all rescued colonies shared mutations in the same gene or genes, while other bypasses showed recurrent mutations in only a subset of colonies (Figure 2B). We then examined the functions of these recurrently mutated genes. For *metA*, *cysE*, and *ilvA*, their annotations suggested plausible mechanisms of rescue, which we investigated further; for the remaining functions, no clear mechanism emerged.

Thus, functional rescue followed diverse genetic trajectories, from adaptation without detectable sequence changes to recurrent mutations shared across independent populations.

### 3. Genetic background and ecological context shape metabolic evolvability

We next examined whether functional-rescue probability and genetic trajectories varied across evolutionary contexts. Our experiments varied two factors associated with spatial structure: genetic background (weak biofilm-forming 168 vs. biofilm model NCIB3610) and initial selective environment (direct MSgg exposure vs. an initial passage on nutrient-gradient plates).

Despite >99% genome sequence similarity between NCIB3610 and 168 (Figure S5), rescue was unevenly distributed: 30 of 46 rescued populations arose in NCIB3610, *hom* and *metA* were bypassed only in this background, and four deletion strains were rescued only after nutrient-gradient initiation. Rescue frequencies ranged from 10.0% in 168 populations initiated directly in MSgg to 33.3% in gradient-initiated NCIB3610 populations (Figure 3A). Genetic background and starting environment were independently associated with rescue probability: The odds of rescue were 2.33-fold higher in NCIB3610 than in 168 (*P*_strain_= 0.015) and 2.27-fold higher after nutrient-gradient than homogeneous MSgg initiation (*P*_environment_ = 0.031; Figure 3B). There was no strain-by-environment interaction (*P*_interaction_ = 0.936).

**Figure 3.**
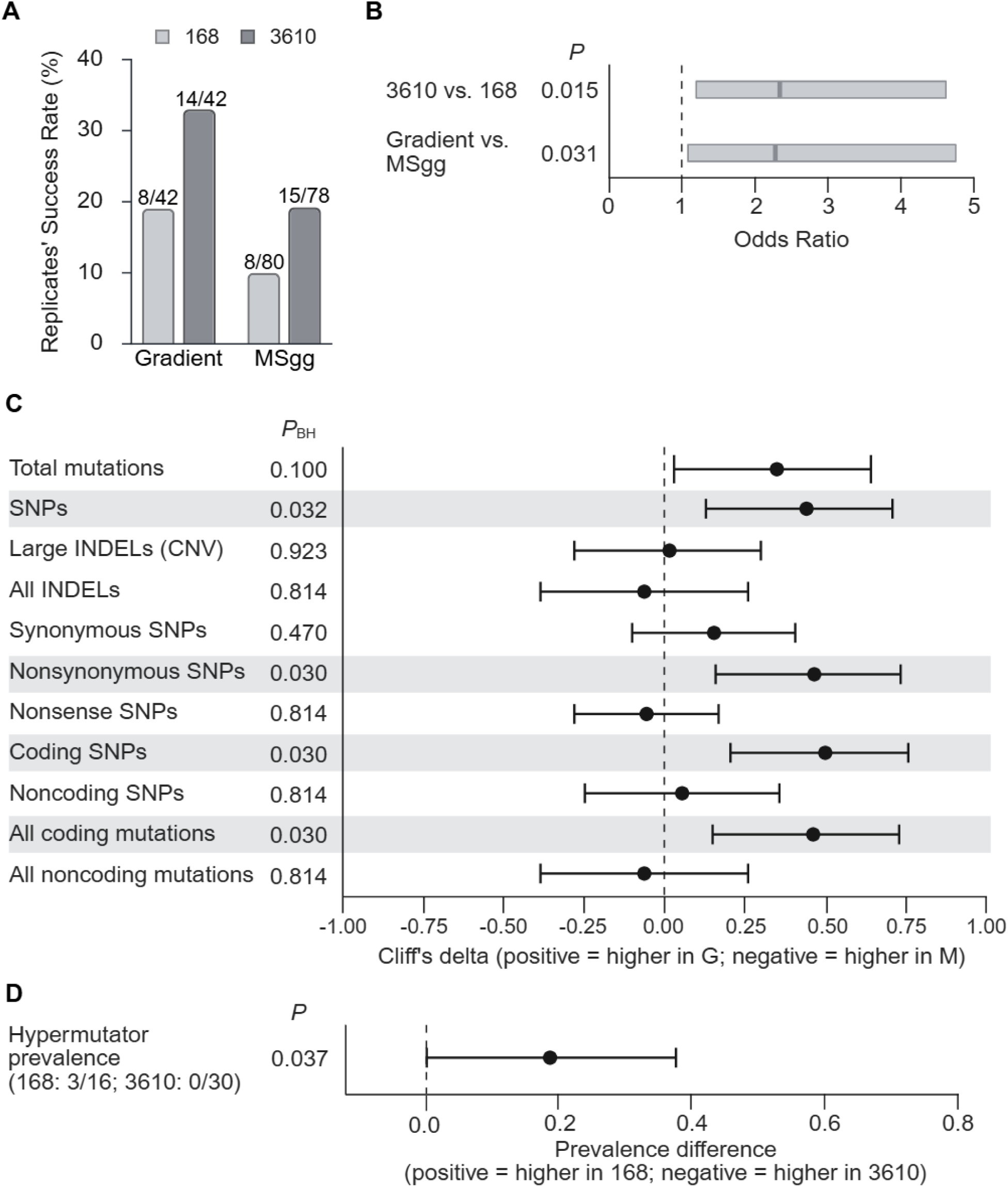
Genetic background and ecological context influenced functional rescue and associated genetic trajectories. **(A)** Functional-rescue frequencies across genetic backgrounds and starting environments. Bars show rescue frequency, with labels indicating successful/initiated populations. **(B)** Main effects of genetic background and starting environment estimated by a binomial generalized estimating-equation (GEE) model clustered by deleted metabolic function. Rectangles indicate 95% confidence intervals, and odds ratios are marked in dark gray; the dashed line at 1 indicates no effect. Genetic background (*P* = 0.015) and starting environment (*P* = 0.031) were significant, but their interaction was not (*P* = 0.936). **(C)** Mutation-category differences across starting environments. Colony-derived counts were summarized at the population level and compared between nutrient-gradient (G) and homogeneous MSgg (M) initiation. Points show Cliff’s delta; positive values indicate higher counts in G and negative values indicate higher counts in M. Lines indicate bootstrap 95% confidence intervals, and gray rows indicate significant categories. Two-sided Mann–Whitney U-test P-values were adjusted across 11 categories using Benjamini–Hochberg correction (*P*_BH_). **(D)** Hypermutator prevalence in the 168 and 3610 backgrounds. Populations were classified as hypermutator-positive when ≥1 sequenced colony was a hypermutator. The point shows the prevalence difference between backgrounds with a bootstrap 95% confidence interval.

The starting environment was also associated with mutation profiles (Table S4). Colonies from gradient-initiated populations had 1.75-fold more total SNPs (2.18 vs. 1.25 mutations per population; Cliff’s delta = 0.440, *P*_BH_ = 0.0316), 2.32-fold more nonsynonymous SNPs (1.48 vs. 0.64; delta = 0.465, *P*_BH_ = 0.0299), 2.03-fold more coding-region SNPs (1.80 vs. 0.89; delta = 0.500, *P*_BH_ = 0.0299), and 1.71-fold more total coding mutations (2.25 vs. 1.32; delta = 0.463, *P*_BH_ = 0.0299) than MSgg-initiated populations(Figure 3C). No mutation category differed significantly between 168 and NCIB3610 after multiple-comparison correction. Hypermutators, however, occurred exclusively in 168: three of 16 populations contained at least one hypermutator colony, compared with none of 30 NCIB3610 populations (*P*_Fisher_ = 0.0369) (Figure 3D). NCIB3610 grew approximately twice as fast as 168 (Figure S6), consistent with previous links between slow growth under stress and hypermutator emergence ^39–41^.

Thus, genetic background and ecological context influenced both access to functional rescue and associated genetic trajectories. Nevertheless, eight functions remained inaccessible under all conditions, indicating that environmental structure increases access to evolutionary solutions without eliminating intrinsic constraints on which metabolic functions can be bypassed.

### 4. Functional rescue reveals alternative routes through central metabolism

To investigate rescue mechanisms and facilitate cross-species comparisons, we characterized mutational convergence associated with *metA*, *cysE*, and *ilvA*.

### Loss of CymR reveals an alternative route for methionine biosynthesis

*metA* encodes homoserine-O-acetyltransferase, which transfers an acetyl group from acetyl-CoA to homoserine, producing O-acetyl-L-homoserine, an essential intermediate in both canonical *B. subtilis* methionine-biosynthesis routes (Table S1) ^42^. Every population and colony bypassing *metA* carried mutations in *cymR*, encoding the global sulfur-metabolism regulator (Table S4) ^43,44^. Three mutations introduced premature stop codons; one was the only mutation detected in three colonies. These Δ*metA* Δ*cymR* deletion strains grew in minimal medium within five to nine days versus three days for wild-type, demonstrating that CymR loss alone restores methionine biosynthesis without MetA (Figure 4A).

**Figure 4.**
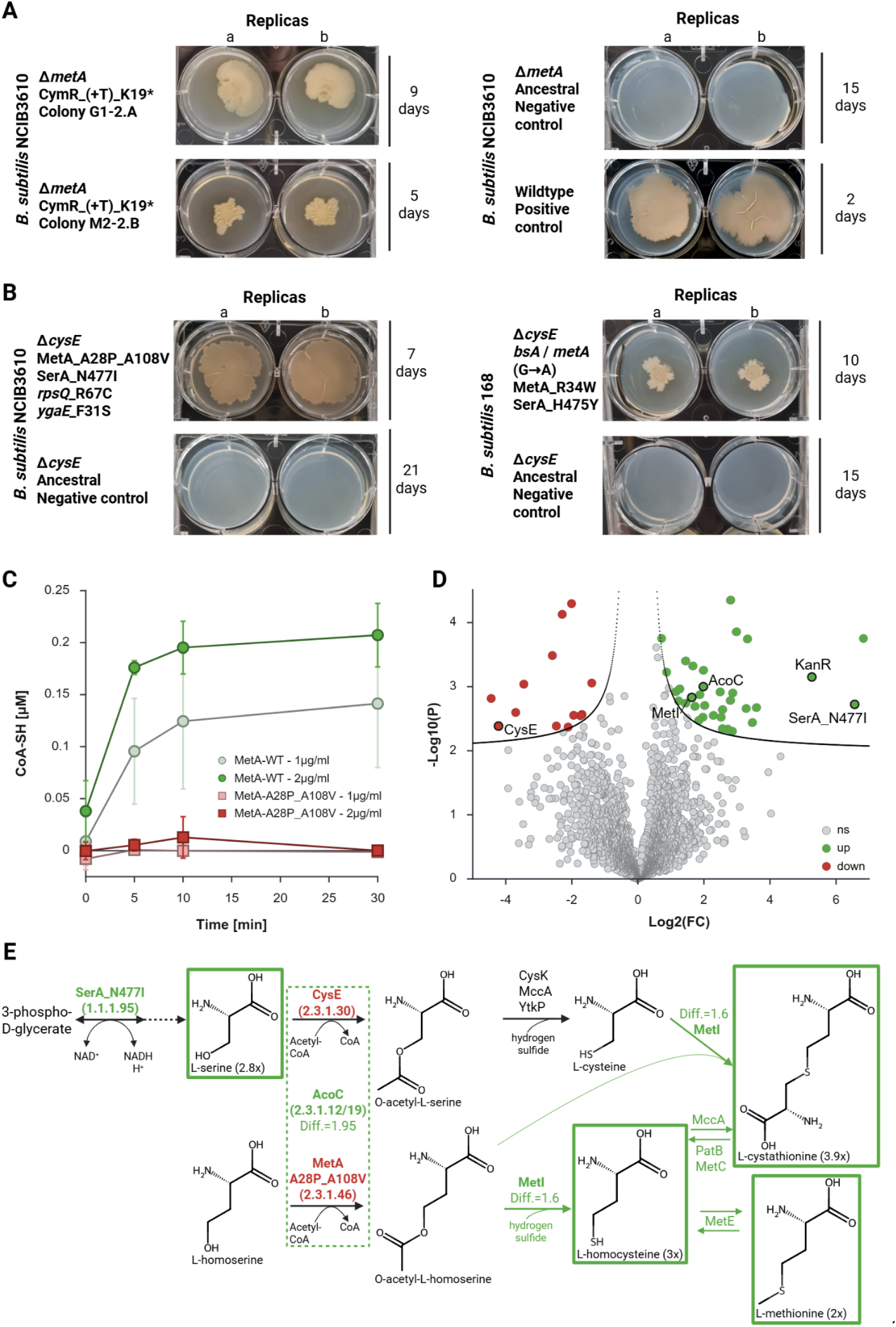
*B. subtilis* Δ*metA* and Δ*cysE* auxotrophs restore methionine and cysteine biosynthesis. **(A)** Selected *B. subtilis* Δ*metA* colonies growing on MSgg agar. Two technical replicates (a, b) of evolved colonies and their ancestor are shown in 6-well plates, with NCIB3610 WT as a reference. Growth time is shown on the right and genotype on the left. **(B)** Selected Δ*cysE* colonies growing on MSgg agar as in (A). **(C)** Enzymatic activity assay of MetA. Homoserine acetyltransferase (MetA) activity was measured as the release of Coenzyme A (CoA-SH), quantified using Ellman’s reagent (DTNB) by measuring TNB²⁻ absorbance at 405 nm, adapted from. Reactions contained homoserine (2 mM), acetyl-CoA (0.4 mM), and purified wild-type *B. subtilis* NCIB3610 MetA (MetA-WT; green) or the MetA-A28P_A108V variant (red) at 1 µg/ml (lighter shades) or 2 µg/ml (darker shades). Measurements were taken at 0, 5, 10, and 30 min. CoA-SH concentrations were calculated from a standard calibration curve (Figure S8). Data represent the mean and the standard deviation of three independent protein purifications (*n* = 3), each measured in three technical replicates. **(D)** Differential proteomic analysis of *B. subtilis* NCIB3610 Δ*cysE*_MetA(A28P/A108V)_SerA(N477I) relative to the wild-type strain. The volcano plot shows changes in protein abundance. The x-axis represents the log₂ fold change, and the y-axis represents the −log₁₀ *P* value. Proteins are classified as significantly increased (upregulated, in green) or decreased (downregulated, in red), whereas nonsignificant proteins are shown in gray. The curved boundaries indicate the significance threshold (FDR = 0.05, *S*₀ = 0.1). Table S8 provides the curve coordinates and raw data. Proteins discussed in the text are annotated: AcoC, acetoin dehydrogenase; CysE, serine acetyltransferase; KanR, kanamycin resistance; MetI, cystathionine gamma-synthase/ acetylhomoserine thiolyase; and SerA_N477I, mutated phosphoglycerate dehydrogenase. **(E)** A simplified schematic representation of the suggested methionine-cysteine biosynthetic pathways bypassing the functions of CysE and MetA. The conventional routes from L-serine and L-homoserine are shown in red, and the proposed bypass in green. Metabolites with increased concentrations are in green frames, and the increase ratio is specified in parentheses.

Because CymR represses 42 sulfur-metabolism genes ^32,44^, we investigated possible routes to MetA bypass. We identified no distant homologs of MetA or the functionally equivalent MetX ^45,46^. However, *B. subtilis* encodes 40 acyl group-transferases (Table S5), one of which could potentially acetylate homoserine promiscuously, although none is regulated by CymR. Alternatively, methionine biosynthesis could proceed via O-phospho-L-homoserine, as in plants by cystathionine γ-synthase (CGS) ^47,48^ and in some *Streptomyces* by homocysteine synthase (MetM, with the sulfur carrier MetO) ^49^.

*B. subtilis* produces O-phospho-L-homoserine during threonine biosynthesis ^50^, and encodes three distant CGS homologs—MetI (36% seq. id.), MetC (37%), and MccB (40%)—plus ThrC (33%), a distant MetM homolog (Table S6). Notably, MccB belongs to the CymR regulon, linking CymR loss to a potential phosphohomoserine-dependent route (Figure S7).

### CysE bypass involves coordinated changes in serine and sulfur metabolism

Populations bypassing *cysE*, encoding serine O-acetyltransferase, and their four colonies carried *metA* and *serA* mutations: MetA(R34W) and SerA(H457Y) in 168, and MetA(A28P/A108V) and SerA(N477I) in NCIB3610. With only one or two additional non-convergent mutations per colony, *metA*/*serA* mutations could suffice for rescue. All colonies grew robustly in minimal medium, with NCIB3610-derived colonies growing faster than 168-derived colonies (Figure 4B), so we further characterized the NCIB3610 mutant.

Because MetA and CysE catalyze analogous acetylations of similar substrates, we hypothesized that evolved MetA acquired CysE activity. Wild-type MetA and A28P/A108V were purified and assayed with MetA and CysE substrates: homoserine and serine. Wild-type MetA retained native activity but showed none with serine, whereas MetA(A28P/A108V) was inactive with both, demonstrating loss of function (Figure 4C).

Despite losing CysE and MetA activity, NCIB3610 *metA*/*serA* mutant retained wild-type-like O-acetyl-serine and O-acetyl-homoserine levels, indicating another source supplies these intermediates. It also showed increased concentrations of serine (2.8-fold, *P* = 0.0006), methionine (2.1-fold, *P* = 0.0057), homocysteine (3-fold, *P* = 0.0436), and cystathionine (3.9-fold, *P* = 0.0021) (Table S7), consistent with greater serine availability and altered sulfur amino-acid flux. Proteomics identified 34 increased proteins (Table S8), including SerA(N477I), consistent with elevated serine concentrations, and O-succinylhomoserine lyase MetI, which produces homocysteine and cystathionine ^42^. AcoC, the dihydrolipoamide acetyltransferase of the acetoin-cleaving system ^51^ and one of the 40 candidate acyltransferases identified above (Figure S7), was also increased in abundance (Figure 4D), although its activity on serine or homoserine remains untested. Thus, CysE bypass may combine increased precursor availability with recruitment of an unidentified acetyltransferase supporting cysteine and methionine biosynthesis (Figure 4E).

### Loss of ThrC promiscuity enables IlvA bypass

Seven populations bypassed *ilvA*, encoding threonine dehydratase, which produces 2-oxobutanoate in isoleucine biosynthesis. Of these, 57% carried mutations in *thrC*, encoding threonine synthase. Three populations and their colonies shared ThrC(T55M), with few additional non-convergent mutations. In one colony, growing within four days, T55M was the only detected mutation, suggesting it alone is sufficient for rescue (Figure 5A).

**Figure 5:**
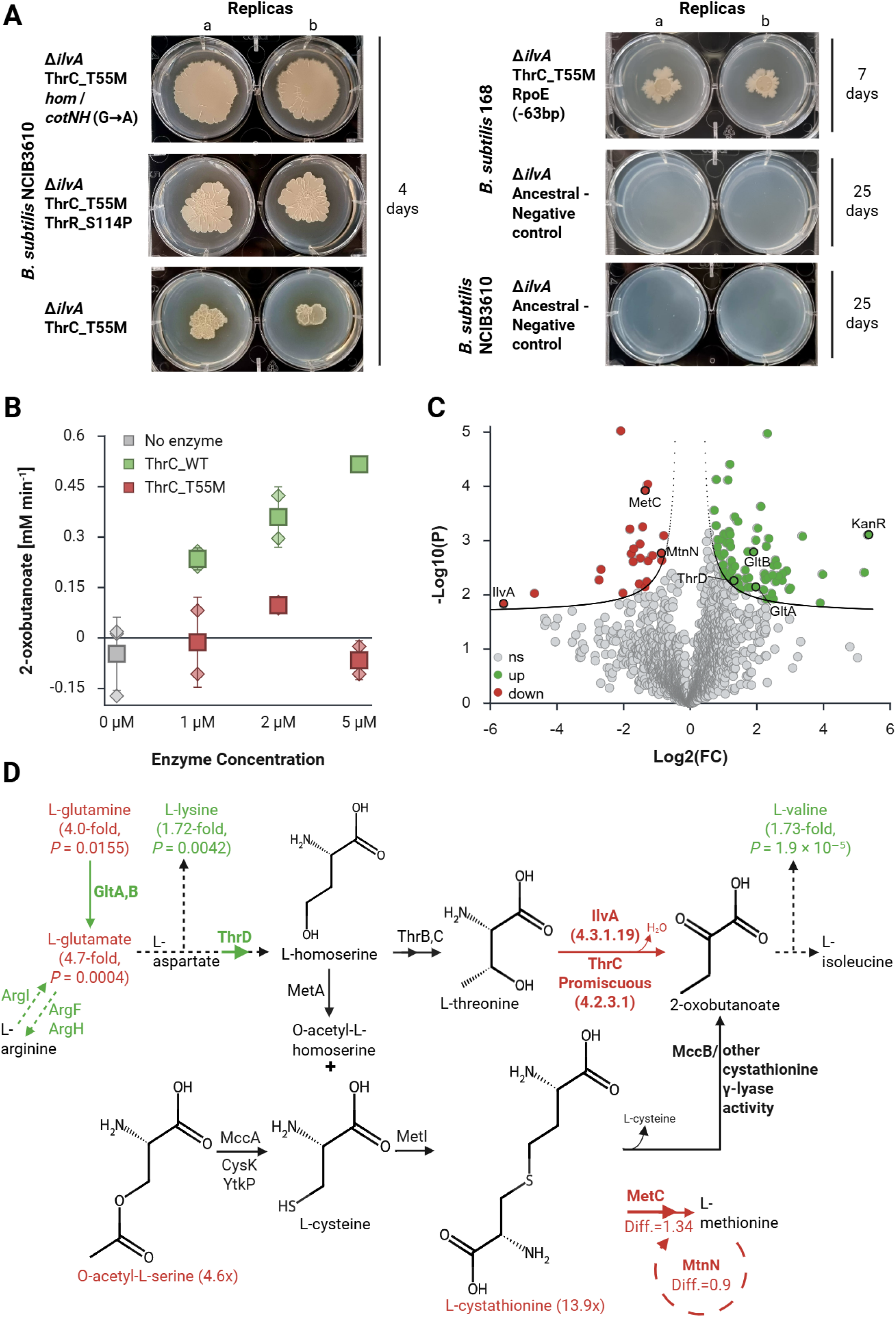
*B. subtilis* Δ*ilvA* auxotrophs redirect isoleucine biosynthesis by silencing ThrC’s promiscuous IlvA function. **(A)** Selected colonies growing on MSgg-Agar. Two technical replicates (a, b) of evolved colonies and their ancestors are shown in 6-well plates. Growth time is shown on the right and genotype on the left. **(B)** ThrC enzymatic activity assay. Threonine dehydratase (IlvA) activity was measured as 2-oxobutanoate production from L-threonine using a DNPH assay at 370 nm, adapted from ^52^. Reactions contained L-threonine (50 mM), PLP (0.5 mM), and purified wild-type *B. subtilis* NCIB3610 ThrC (ThrC-WT; green) or ThrC-T55M (red) at 0 (No enzyme; gray), 1, 2, or 5 µg/mL. Measurements were taken at 0, 5, 10, 30, and 60 min. 2-Oxobutanoate concentrations were calculated from a standard curve (Figure S9), and activity is shown as its production rate. Squares show means, and diamonds indicate individual values from two replicate experiments (*n* = 2). **(C)** Differential proteomic analysis of *B. subtilis* NCIB3610 Δ*ilvA*_ThrC(T55M) relative to the wild-type strain. The volcano plot shows changes in protein abundance. The x-axis represents the log₂ fold change, and the y-axis represents the −log₁₀ *P* value. Proteins are classified as significantly increased (upregulated, in green) or decreased (downregulated, in red), whereas nonsignificant proteins are shown in gray. The curved boundaries indicate the significance threshold (FDR = 0.05, *S*₀ = 0.1). The curve coordinates and raw data are provided in Table S8. Proteins discussed in the text are annotated: GltA/B, glutamate synthase; IlvA, threonine dehydratase; KanR, kanamycin resistance; MetC, cystathionine beta-lyase; MtnN, methylthioadenosine nucleosidase; and ThrD, aspartokinase III. **(D)** A simplified schematic representation of the suggested isoleucine biosynthetic pathways bypassing the IlvA function of IlvA and promiscuous ThrC. Red arrows represent the conventional route from threonine to 2-oxobutanoate and reactions of downregulated enzymes– MetC and MtnN (with |log2FC| values); green arrows represent reactions of upregulated enzymes–GltA, GltB, ArgF, ArgH, ArgI, and ThrD. Metabolites with decreased concentrations are in red, and those with increased concentrations are in green, with the change ratio in parentheses. The light green background follows the proposed bypass from acetyl serine to 2-oxobutanoate, including the broader changes from glutamine.

Because wild-type ThrC has weak promiscuous IlvA activity ^50^, we hypothesized that T55M enhanced it and replaced IlvA. Instead, T55M abolished this activity and was predicted to strongly destabilize ThrC (ΔΔG = 4.32), excluding enhanced promiscuous 2-oxobutanoate production as the mechanism (Figure 5B). We therefore compared wild-type and evolved metabolomes and proteomes. ThrD, which converts aspartate semialdehyde to homoserine, was upregulated. Because homoserine branches toward either the canonical ThrC/IlvA pathway to 2-oxobutanoate or sulfur amino-acid metabolism (Figure 5D), increased ThrD could supply both. The evolved *ilvA* deletion strain was also depleted of O-acetyl-L-serine (4.7-fold, *P* = 0.0004) and cystathionine (13.9-fold, *P* = 0.038) (Table S7), intermediates of cysteine and methionine metabolism ^42^. *B. subtilis* can produce 2-oxobutanoate from cystathionine through cystathionine γ-lyase activity, including MccB ^42^. MetC, which directs cystathionine toward homocysteine and methionine, and MtnN, a methionine-salvage component, were downregulated, indicating altered methionine/SAM metabolism. These changes support increased homoserine supply and redistribution of sulfur-metabolism intermediates toward 2-oxobutanoate production.

Broader amino-acid remodeling accompanied this shift. Glutamine and glutamate were depleted (4.0-fold, *P* = 0.0155; 4.7-fold, *P* = 0.0004), while glutamate synthase GltAB was upregulated (Table S8), consistent with altered glutamate homeostasis. Several arginine-metabolism enzymes were also upregulated, whereas lysine, an aspartate-derived amino acid, accumulated (1.72-fold, *P* = 0.0042). Valine likewise accumulated (1.73-fold, *P* = 1.9 × 10⁻⁵), indicating additional branched-chain amino-acid metabolism remodeling. Thus, restoring 2-oxobutanoate reconfigures interconnected pathways to redistribute metabolic flux while preserving other amino-acid branch outputs (Figure 5D).

Together, these results show that functional rescue can arise through loss-of-function mutations that redirect metabolic flux through latent pathways rather than optimize a direct replacement enzyme.

### 5. Evolutionary routes to functional rescue differ between *B. subtilis* and *E. coli*

Finally, we asked whether *B. subtilis* rescue patterns reflect general bacterial-network properties or organism-specific solutions. The closest experimental analog is Blank et al. ALE study in *E. coli* ^13^, which evolved five replicates of 87 auxotrophic strains (435 populations) in liquid minimal medium, initially forming a nutrient gradient, enabling slow adaptive recovery and identification of compensatory metabolic innovations. Overall, 68/435 populations (16%) regained growth, rescuing 22/87 functions (25%).

Although overall recovery was similar (16% in *E. coli*; 19% in *B. subtilis*), *B. subtilis* bypassed more tested functions (9/17, 53%) vs. 22/87 (25%) in *E. coli*. Among 12 auxotrophic functions tested in both species, *B. subtilis* bypassed seven (*argH, aroC, cysE, ilvA, leuB, pheA,* and *serB*) and *E. coli* only two (*ilvA* and *serB*) (Figure 6A). Gene-level convergence was also lower in *B. subtilis*: of 22 bypassed *E. coli* functions, 16 were rescued in multiple independent populations, and 10 (62.5%) had the same gene mutated in every rescuing population, versus 3 out of 9 *B. subtilis* functions (33%). The number of successful populations differed among functions (Figure 2B). Thus, *B. subtilis* populations more frequently recruited different genes to overcome the same lesion.

**Figure 6:**
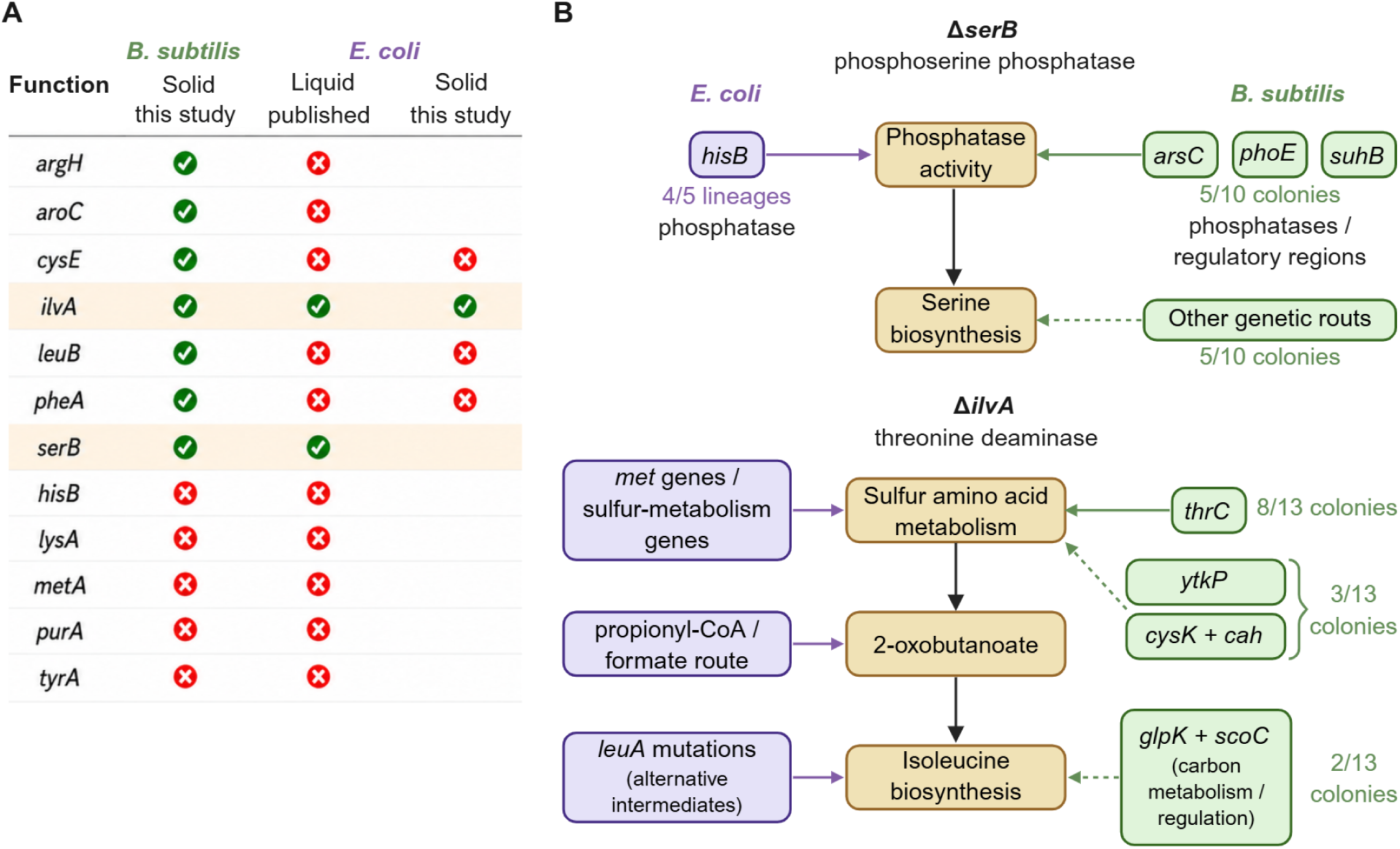
Functional rescue differs across species despite convergence on specific metabolic solutions. **(A)** Functional bypassability of auxotrophies tested in both *B. subtilis* and *E. coli* under previously published liquid-culture conditions ^13^ and under the structured solid-medium conditions used in this study. Green checks indicate bypass; red crosses indicate no bypass. **(B)** Genetic and metabolic routes associated with Δ*serB* and Δ*ilvA* bypass in both species. Solid arrows indicate experimentally supported or previously demonstrated connections; dashed arrows indicate putative pathway associations.

We next examined *ilvA* and *serB*, the two functions bypassed in both species, for pathway-level similarities despite gene-level divergence. After losing *serB*, four of five *E. coli* lineages accumulated mutations in the phosphatase gene *hisB* ^13^ were also implicated in an independent study ^53^. Instead, five of ten *B. subtilis colonies* mutated phosphatases or their regulatory regions (ArsC, SuhB, or broad-substrate-specificity PhoE ^54^), while the other half lacked phosphatase-related mutations, indicating additional rescue routes. Thus, both species recruited phosphatase-related functions through different genes (Figure 6, Table S4), and in *B. subtilis* this was not the only solution.

*ilvA* loss produced multiple solutions in both species. Three *E. coli* studies identified *met,* or other sulfur-metabolism mutations ^13–15^, suggesting the rerouting of methionine-biosynthesis intermediates toward 2-oxobutanoate production ^14^. Additional solutions include a propionyl-CoA/formate-dependent route ^14^, and recurrent *leuA* mutations that may restore isoleucine biosynthesis through an unknown mechanism 15. Like *E. coli*, *B. subtilis*’ ThrC(T55M) restored 2-oxobutanoate through redirecting sulfur amino-acid metabolism (Figure 5). Other mutations, like in *glpK*/*scoC*, suggest altered carbon metabolism and regulation, whereas *ytkP*/*cysK* are associated with cysteine biosynthesis and may represent additional sulfur-metabolism routes. Thus, both species access multiple solutions to IlvA loss, with sulfur metabolism repeatedly emerging as pathway-rather than gene-level convergence (Figure 6).

These differences could reflect metabolic-network architecture or evolutionary environment. Indeed, selective conditions alter Δ*ilvA* solutions ^15^, and physical structure, including biofilms, can alter evolutionary trajectories ^55,56^. Previous *E. coli* experiments were predominantly in liquid culture ^13–15^, whereas our *B. subtilis* populations evolved on solid media with spatial structure, local nutrient gradients, and sometimes biofilms. To test whether these environmental differences could explain the species-specific rescue patterns, we experimentally evolved *E. coli* Δ*cysE*, Δ*ilvA*, Δ*leuB*, and Δ*pheA* under the same structured conditions used for *B. subtilis*. Our *E. coli* strains did not form biofilms, preventing a direct comparison of biofilm effects. Only IlvA was bypassed, matching previous liquid-selection results (Figure 6). Thus, for these four functions, spatial structure did not reproduce *B. subtilis’* rescue capacity, suggesting species-specific metabolic networks define available routes, while the environment influences their accessibility.

## Discussion

Here, we explored the capacity of *Bacillus subtilis* to evolve after the loss of 17 biosynthetic functions across two genetic backgrounds and starting environments. Nine functions were bypassed, whereas eight resisted evolution. Rescue was more likely in the NCIB3610 background and after nutrient-gradient initiation, which was also associated with altered mutation profiles. Thus, rescue depends on both the metabolic lesion and ecological context.

That only a subset of functions could be bypassed indicates substantial but constrained metabolic plasticity. This is consistent with previous work showing that the ability of latent or redundant activities to compensate for gene loss depends on their connectivity within the metabolic network and on regulatory and physiological context ^57–60^. Experimental flux analysis in *B. subtilis* likewise revealed limited redistribution following gene deletion ^61^. Together, these studies suggest that the presence of potentially redundant enzymes alone should be insufficient to predict functional rescue. Consistent with this expectation, neither paralogue number nor the number of enzymes with related EC classifications predicted bypassability in our experiments.

Comparison with *E. coli* further suggests that these constraints are species-dependent. Despite similar population recovery frequencies, a larger fraction of functions was bypassed in *B. subtilis* than in *E. coli.* The *E. coli* screen tested about five times as many auxotrophic functions as those examined here, and only 12 were shared. Thus, differences in bypassed functions may partly reflect the sample size and auxotrophy type. Despite these differences, *serB* and *ilvA* revealed biochemical convergence: different genes were recruited, but phosphatase-related functions repeatedly appeared during SerB rescue ^13,53,54^, and sulfur amino-acid metabolism during IlvA bypass ^13–15^. Thus, genetic routes appear organism-specific even when evolution accesses related biochemical strategies.

Ecological conditions also influenced which solutions *B. subtilis* reached. Nutrient-gradient initiation increased rescue and altered mutation profiles, consistent with evidence that selective conditions can change how the same metabolic lesion is compensated ^15^. Yet spatial structure did not reproduce broader *B. subtilis* bypassability in selected *E. coli* auxotrophs. Environmental context therefore affects route accessibility, while metabolic network architecture constrains the repertoire of potential solutions.

The three mechanistically characterized bypasses—Δ*metA*, Δ*cysE*, and Δ*ilvA*—show that metabolic innovation can result from reducing or losing existing activities rather than enhancing a compensating enzyme. Previous bacterial studies have shown that removing metabolic or regulatory constraints can expose latent flux routes without creating new biochemical capabilities ^62,63^. Functional innovation can therefore arise not only by enhancing underground activity, but also by eliminating activities that constrain alternative uses of the existing network. This is particularly striking for Δ*ilvA*. Previous *B. subtilis* Δ*ilvA* revertants selected with threonine restored growth by derepressing the *hom-thrCB* operon and exploiting the ThrC’s promiscuous threonine-dehydratase activity ^62^. In contrast, our populations evolved without threonine, and the recurrent ThrC(T55M) mutation abolished the promiscuous activity while enabling rescue through sulfur amino-acid metabolism. These opposing outcomes illustrate how the environment can determine which latent metabolic solution becomes advantageous.

Overall, metabolic evolvability reflects an interaction between network architecture and evolutionary context. Network organization constrains available compensatory routes, while environmental conditions influence their accessibility. Differences between *B*.

*subtilis* and *E. coli*, together with occasional pathway-level convergence, show that latent metabolic solutions cannot be extrapolated across organisms. Exploring diverse metabolic networks should reveal additional forms of functional rescue and broader principles of metabolic evolution.

## Materials and Methods

### Bacterial growth and maintenance

Table S9 lists the bacterial strains and plasmids used in this study. *B. subtilis* was grown and maintained in Luria-Bertani (LB) (Bactoᵀᴹ) medium with Kanamycin (Kan, 50 μg/mL) (Tivan Biothec) if it was one of the auxotrophic strains or evolved from them. For solid media, we used LB plus 1% agar. To test for auxotrophy or long-term incubations to promote adaptation, the chemically defined solid minimal medium (MSgg) was used, containing 5 mM potassium phosphate, 100 mM MOPS (pH 7.5), 2 mM MgCl_2_, 50 µM MnCl_2_, 1 µM ZnCl_2_, 2 µM thiamine, 0.5% glycerol, 0.5% glutamate, 700 µM CaCl_2_, 50 µM FeCl_3_; adapted from ^64^. We solidified it by adding Noble agar (Difco) to a final concentration of 1.5%. The medium was supplemented with 50 µg/mL L-tryptophan for *B. subtilis* 168.

### Strain isolations and storage

The strain collection was indexed in a 96-well plate as frozen glycerol stocks. Each mutant was streaked on LB agar + Kan and incubated overnight at 37°C. Four isolated colonies were collected and grown in 3 mL LB-Kan at 37°C overnight (O/N) with agitation. 800 μL from each culture was stored in 30% glycerol at-80°C. The remaining culture was used to extract DNA using the GenEluteᵀᴹ Bacterial Genomic DNA Kit.

### Genotypic validation

The expected genotype (gene deletion) was verified by conventional (qualitative) PCR using primers designed with Primer3 ^65^ (Table S9). One set of primers amplified a small fragment (300–400 bp) of the deleted gene, and another set amplified the Kan resistance cassette (1 Kbp) at its specific insertion site, using the unique flanking regions of each knocked-out gene ^22^.

### Bacillus subtilis NCIB3610 transformation

Genetically engineered *B. subtilis* NCIB3610 strains were obtained by natural competence, as previously described ^64^. A *B. subtilis* NCIB3610 wild-type colony was cultured in 1 mL of media, prepared by diluting in water 10X of MC (0.5% glucose, 1.4% K_2_HPO_4_, 0.6% KH_2_PO_4_, 30 nM sodium citrate tribasic dihydrate, casein hydrolysate, ammonium iron (III) citrate, L-glutamic acid potassium salt monohydrate, and 3 μM MgSO_4_) enriched with 3 μL 1 M MgSO_4_, 10 μL 0.5% tryptophan, and 10 μL 0.5% histidine, adapted from ^66^. It was then incubated for 5 hours at 37°C at 150 rpm. 1 μg of genomic DNA from each chosen *B. subtilis* 168 mutant was incubated with 500 μL of culture for one hour under the same conditions. The culture was resuspended in 100 μL and plated on LB-agar+Kan. Single colonies of *B. subtilis* NCIB3610 mutants were purified and genotyped as described before.

### Nutrient gradient environment (’Gradient-plate’) calibration and preparation

Standard Petri dishes (diameter 90 mm) were filled with 40 mL agar-MSgg. Once solidified, a 2 cm-diameter cavity was made in the center, and 1-1.5 mL of LB-agar (1.5%) was gently loaded using a pipette. Next, we calibrated the optimal gradient starting point by changing the inoculation time after plate preparation. We assumed that the longer the time elapsed after plate preparation, the more nutrients from the LB center would diffuse into the minimal medium, thereby exerting different selective pressures. We thought to start with the least permissive. In this, we inoculated the LB center with several auxotrophic strains immediately after plate preparation, or after 24 or 48 hours, when the plates were kept sterile at room temperature. After inoculation, the plates were sealed with parafilm, incubated at 30°C, and monitored daily.

We observed rapid, homogeneous growth in plates left to form a gradient for 24 or 48 hours, as the auxotrophic strains took over the plate within 2-3 days. However, the same strains had greater difficulty growing on freshly made plates, indicating that they exerted selection pressure (Figure S1). Thus, using this setup, we tested the 34 strains shown herein.

### Auxotroph evolution

A single colony from the frozen glycerol stock was isolated on an LB-agar+Kan plate O/N at 37°C. The colony was incubated in 2 mL LB at 37°C with shaking until the OD600 reached 0.6-0.8, and 5 μL was used to inoculate a gradient plate. The remaining culture was washed 4 times in MSgg without glycerol or glutamate (MS), and 5 μL of the washed cells was inoculated into a single agar-MSgg well (6-well plate, one strain per plate). Both plates were sealed with parafilm, incubated at 30°C with the plates facing up, and monitored daily.

After simultaneous inoculation onto the nutrient gradient and MSgg minimal medium plates, the plates were incubated at 30°C, with the plates facing up, for up to 3 months. Growth arrest was simultaneous and determined by the population growing on the gradient once it covered most of the plate surface. We collected and homogenized the entire biomass from each plate or well in MS solution as a single population. Part of each sample was reserved for genomic DNA extraction and genotypic validation as before, and the rest was resuspended in 30% glycerol for long-term storage at-80°C. To start the following growth cycle, a population stock was slightly thawed, and 20 μL was taken to inoculate 3 mL LB-Kan O/N. From this culture, 2 mL of LB was inoculated to obtain OD600 = 0.01-0.02 (5-10μL) and grown to reach a final OD600 = 0.6-0.8. As in the previous step, cells were washed before inoculating new MSgg 6-well plates. Once grown, the biomass was again collected as a population, homogenized, genotyped, and stored before the next passage.

### Colony isolation from evolved populations

Single-colony isolation was performed by streaking an agar-MSgg plate directly from the glycerol stock of the last evolved population and incubating at 30°C. After growth, four colonies were picked to re-streak fresh agar-MSgg plates, ensuring independent growth on minimal media. A single colony from each plate was then cultured overnight in 3 mL LB-Kan (50 μg/mL) at 30°C for DNA extraction, glycerol stock, and genotyping. We sequenced and analyzed two colonies per population.

### Quantifying growth rate differences between *B. subtilis* NCIB3610 and 168 on MSgg agar plates

Overnight cultures were initiated by inoculating 3 mL of LB and incubating at 30°C and 250 rpm. The next day, cultures were refreshed (1:30 into 3 mL LB) and grown to an OD600 of 0.3–0.8. Cells were washed three times (3,200 rcf, 5 min) with 1 mL MSgg to remove residual LB and resuspended in MSgg, then normalized to OD600 = 0.01 in 500 µl. 5 µl were spotted at the center of wells containing 3 mL MSgg agar (in 12-well plates). For CFU enumeration, biofilm material from each well was collected with a sterile loop into 3 mL saline in a 15 mL tube, vortexed for 10 seconds to homogenize, and serially diluted in saline. 100 µL of appropriate dilutions was spread on LB agar using glass beads, followed by overnight incubation at 30°C. At later time points, suspensions were adjusted to OD600 = 0.1 in 450 µL in a deep-well plate and then serially diluted before plating.

### Identification of adaptive suppressor mutations

Ancestral clones, evolved populations, and selected isolated colonies from each auxotrophic strain were analyzed as follows:

### Whole-genome sequencing

Genomic DNA samples were prepared for Illumina Whole Genome Sequencing (NGS) performed by SeqCenter in Pittsburgh, PA. Sample preparation included DNA extraction, genotyping, and dsDNA quantification by microplate reader at 260 nm. Paired-end sequencing libraries were generated using the Illumina 300-cycle kit (Illumina NextSeq 2000, producing 2×151 bp reads) with the SeqCenter 400 Mbp sequencing package, with a minimum of 400 Mbp and a quality score of Q30 (99.9% accuracy) or higher. Raw reads were provided as FASTQ files, along with sequencing statistics (total read pairs, total reads, total bp > Q30, and % bp > Q30).

### Computational analysis

All reads underwent a quality check using FastQC ^67^ and were trimmed to remove adapters (NexteraPE-PE.fa) using Trimmomatic ^68^. Matching annotated reference genomes were downloaded from NCBI in GenBank files: *Bacillus subtilis* 168 [tax. ID 224308, accession NC_000964] and *Bacillus subtilis* NCIB3610 [tax. ID 535026, accession NZ_CP020102]. RAST application ^69^ was used to re-annotate both references to obtain an accurate alignment. Variant calling was applied by Barrick’s lab Breseq algorithm ^70^, installed and executed as instructed on http://barricklab.org/breseq. Reads were mapped against their matching-strain genome reference – 168 or 3610 – first against the original NCBI reference and then against the RAST-reannotated version (Table S4 shows all mutations’ descriptions and analyses). A second mutation prediction analysis was executed as a comparison using the Snippy tool (Galaxy Version 4.6.0 T, S. (2015)) (Table S4). Variations in read coverage depth of single colonies were obtained through breseq BAM2COV. This additional subcommand was executed after the main pipeline was run using the BAM database files of read alignments to create coverage plots and tables (Table S4 with copy number variation analysis; plots in File S1).

### Statistical analysis of functional rescue and mutation profiles

Functional-rescue success was treated as a binary outcome for each replicate population, with success defined as completion of two consecutive passages in MSgg. The effects of genetic background (168 or NCIB3610) and starting environment (nutrient gradient (G) or homogeneous MSgg (M)) on rescue probability were analyzed using a binomial generalized estimating-equation (GEE) model with a logit link. Deleted metabolic function was used as the clustering variable to account for repeated testing of the same function across experimental conditions. A strain-by-environment interaction was first included to test whether the effect of either factor depended on the other; because no interaction was detected, the main effects of strain and environment were estimated using an additive model. Model coefficients were expressed as odds ratios (OR) with 95% confidence intervals.

Mutation-profile analyses were based on whole-genome sequencing of evolved colonies. To avoid treating colonies isolated from the same evolved population as independent replicates, mutation counts from paired colonies were averaged and the evolved population was used as the statistical unit. Populations in which both sequenced colonies were classified as hypermutators (aroC_168_G_1-2 and pheA_168_M_3-2) were excluded from mutation-count analyses. For ilvA_168_M_1-2, only the non-hypermutator colony was retained, and for the longitudinally sampled leuB_3610_G_2 lineage, only the later leuB_3610_G_2-3.A colony was included. Mutation counts were compared between gradient-and MSgg-initiated populations and between the 168 and NCIB3610 backgrounds using two-sided Mann–Whitney U tests. Cliff’s delta was calculated as a rank-based effect size, with percentile-bootstrap 95% confidence intervals obtained from 10,000 resamples. Within each comparison, P values were adjusted across the 11 mutation-count categories using the Benjamini–Hochberg procedure to control the false-discovery rate and are reported as (*P*_BH_). Hypermutator prevalence was analyzed separately: a population was classified as hypermutator-positive when at least one sequenced colony was identified as a hypermutator, and prevalence between the 168 and NCIB3610 backgrounds was compared using a two-sided Fisher’s exact test.

### Plasmid construction for protein production

The DNA coding sequence, either ThrC or MetA, was amplified by PCR using the isolated *B. subtilis* genomic DNA sequence as template and the corresponding primers (Table S9). The plasmid backbone was amplified by PCR from an empty pET28a(+) vector containing a StrepTag affinity tag on the C-terminus of the multiple cloning site. The amplification reactions were performed using Q5^®^ polymerase according to the manufacturer’s instructions (New England Biolabs). Then, the two previously amplified fragments were verified by standard gel electrophoresis, purified from the reaction using the QIAquick PCR purification kit (QIAGEN Labs, US), and assembled into a single construct using the NEBuilder® HiFi DNA assembly reaction, as indicated by the manufacturer (New England Biolabs). The construct was then transformed into NEB 5-alpha competent *E. coli*, a derivative of DH5α (New England Biolabs), plated on LB agar-kan, and incubated overnight at 37°C. The functional plasmids were extracted from overnight cultures of the positive colonies using the QIAprep spin miniprep kit (QIAGEN Labs) and further sequenced for confirmation. Whole-plasmid sequencing was performed by Plasmidsaurs using Oxford Nanopore Technology, with custom analysis and annotation.

### Protein expression and purification (MetA, ThrC, and variants)

The constructed and verified plasmids were transformed into One Shot™ BL21 Star™ (DE3) chemically competent *E. coli* cells (Thermo Fisher Scientific), plated on LB-Kan agar (kanamycin 50 μg/mL), and incubated overnight at 37°C. Transformants were cultured in 5 mL LB-Kan (50 *μ*g/mL) and incubated overnight at 37°C. A 1:20 dilution was then inoculated into 500 mL of LB-Kan (50 μg/mL) and incubated at 37°C until reaching log-phase (OD600 = 0.4-0.8), then induced with IPTG at a final concentration of 0.5 mM and incubated overnight at 18°C. Cells were harvested by centrifugation at 15,400 x g for 20 minutes at 4 °C. The supernatant was discarded.

For ThrC and its variants, the cell pellets were resuspended in 10 mL of 50 mM Tris-HCl buffer (pH 7.5) supplemented with Benzonase® Nuclease (Merck Millipore, Cat# 70746), EZBlock™ Protease Inhibitor Cocktail (EDTA-Free) (BioVision), and chicken egg white lysozyme (Sigma-Aldrich) (1 mg/mL). The cells were lysed by incubation on ice for 20 minutes, followed by sonication for 10 minutes (5-second pulse on, 5-second pulse off, 40% of maximum power). Cell debris was removed by centrifugation at 15,400 x g for 20 minutes at 4 °C. For MetA and its variants, the cell pellets were resuspended in 5 mL of CelLytic™ B Cell Lysis Buffer Benzonase® Nuclease (Merck Millipore, Cat# 70746), EZBlock™ Protease Inhibitor Cocktail (EDTA-Free) (BioVision). The cell suspension was vortexed for 10 minutes for lysis and centrifuged at 15,400 x g for 20 minutes at 4°C.

Following lysis, the clear lysate containing all produced proteins (MetA and its variants, and ThrC and its variants) was loaded onto the corresponding gravity columns containing streptavidin StrepTag® resin (IBA Lifesciences GmbH) for affinity chromatography. The column was washed with 6 column volumes of washing buffer (100 mM Tris-HCl, pH 8, 150 mM NaCl, 1 mM EDTA), and elution was performed in 6 steps of 0.5 column volumes each, using washing buffer supplemented with 25 mM desthiobiotin. The purity of the eluted fractions was evaluated by electrophoresis using a 12.5% polyacrylamide gel. Total protein content was determined using the Coomassie Blue assay with bovine serum albumin as the standard.

### Serine and homoserine acetyltransferase *in vitro* function (CysE and MetA activity)

Acetyltransferase activity was quantified by measuring the release of Coenzyme A (CoA-SH) from acetyl-CoA, adapted to a 96-well plate format. The colorimetric assay is based on the disulfide exchange of CoA-SH with Ellman’s reagent (DNTB 3 mM, EDTA 2 mM), which produces a yellow chromophore, 2-nitro-5-thiobenzoate (TNB⁻), with a maximum absorption wavelength of 405 nm. Reactions (120 μL) contained purified MetA (WT or variant; 1-2 *μ*g/mL), acetyl-CoA (0.4 mM), MgCl₂ (5 mM), and L-homoserine (2 mM) in 50 mM Tris–HCl, pH 7.5, and were incubated at 37 °C for 60 min. Reaction aliquots were quenched by adding guanidine-HCl to 0.8 M (final). A 20 *μ*L aliquot was mixed with 100 μL of the DTNB/EDTA solution, brought to 200 μL with the same assay buffer, incubated for 10 min at room temperature, and read at the specified wavelength. CoA-SH was quantified from a standard curve (Figure S8).

### Threonine dehydratase *in vitro* function (ilvA activity)

Threonine dehydratase activity was assessed by quantifying the 2-oxobutanoate generated by the deamination of L-threonine. The 2-oxobutanoate reacts with 2,4-dinitrophenylhydrazine (DNPH) to form its colored hydrazone; upon alkalinization, the chromophore develops and is read at 370 nm ^52^. Reactions (300 µL) were prepared in 50 mM Tris–HCl, pH 7.5, containing L-threonine (50 mM), PLP (0.5 mM), and purified

ThrC or variants (1, 2, and 5 µM), and incubated at 37 °C for 60 min. At the selected time point, a 40 µL aliquot was transferred to a microplate well and immediately quenched with 10 µL HCl. DNPH derivatization was initiated by adding 15 µL of 0.1% (w/v) DNPH and incubating for 5 min at room temperature. Absolute ethanol (35 µL) was then added and mixed, followed by 85 µL of 2.5 M NaOH; the mixture was incubated for 10 min at room temperature to complete color development. The volume was brought to 200 µL with 15 µL assay buffer, and A_370_ was recorded. A 2-oxobutanoate standard curve was used for quantification (Figure S9).

### Metabolites extraction and metabolomics

Basal metabolites were extracted from exponentially growing *B. subtilis* NCIB3610 wild-type, Δ*ilvA*_ThrC(T55M), and Δ*cysE*_MetA(A28P/A108V) strains using the hot-water extraction method, as follows:

### Chemicals

LC-MS grade acetonitrile (ACN) and LC-MS grade water (H_2_O) were acquired from Honeywell (Charlotte, North Carolina, US), LC-MS grade formic acid (HCOOH) was acquired from Sigma Aldrich (Søborg, Denmark) while LC-MS grade methanol (MeOH) was purchased from Fisher Chemical (Waltham, Massachusetts, USA).

L-serine, L-Isoleucine, L-cysteine, L-Methionine, L-alanine, L-aspartic acid, L-glutamine, and L-arginine all analytical standard grade (98% purity) were acquired from Sigma Aldrich (Søborg, Denmark). Individual 5mM stock solutions of these compounds were prepared in Milli-Q water. The stock solutions were then used to prepare a 10 µM mixture 1 (Mix1) in 50% MeOH in Milli-Q water. This mix was used for quality control purposes for the non-targeted metabolomics analysis.

### Sample preparation

Single colonies were inoculated in 3 mL LB medium (with added kanamycin for mutants) and grown overnight at 37°C. Pre-cultures were washed three times with 1 mL 1xMSgg medium lacking CaCl₂ and FeCl₃, then resuspended in 3 mL of the same medium. The washed pre-cultures were used to inoculate the main cultures (4 mL complete 1xMSgg, no kanamycin) at an initial OD₆₀₀ of 0.01–0.02, which were incubated at 30°C, 200 rpm until OD₆₀₀ ≈ 0.8. Three biological replicates per strain were harvested by centrifugation (6000 × g, 10 min), and metabolites were extracted by resuspending pellets in boiling water (100°C) with two sequential incubations (15 min each, 500 rpm shaking). Supernatants from both extractions were combined, chilled on ice, clarified by centrifugation (12,000 rpm, 5–10 min), and evaporated to dryness in a vacuum concentrator at 30°C.

### Non-targeted LC-MS/MS metabolomics analysis

Semi-quantitative LC-MS/MS metabolomics analysis was performed using a Vanquish Duo UHPLC binary system coupled to an Orbitrap IDXTM mass spectrometer (Thermo Fisher Scientific Co., Waltham, Massachusetts, USA). The chromatographic conditions used were as previously described. In brief, separation was achieved using a ACQUITY UPLC BEH Amide (100 mm × 2.1 mm, 1.7 µm particle size) column (WatersTM, Milford, MA, USA) equipped with an ACQUITY UPLC BEH amide guard (Waters^TM^). The mobile phase consisted of H_2_O + 0.1% HCOOH (A) and ACN + 0.1% HCOOH (B). The following gradient elution was used at a flow rate of 0.35 mL min–1: 0-0.8 min 85%, 0.8-3.2 min 85% to 50% B, 3.2-4.2 min 50% B, 4.2-5.2 min 50% to 30% B. The column was re-equilibrated for 3 min at 85% B. The column was kept at 40°C throughout the analysis. The autosampler temperature was kept at 7 °C, and the injection volume was 1 µL.

The MS/MS data acquisition was performed in positive and negative ion mode using heated electrospray ionization (HESI) with voltages of 3500 V and 2500 V, respectively. Full MS/MS spectra (data-dependent acquisition-driven MS/MS) were acquired using *m/z* range of 70–1000 and in profile mode. The MS1 resolution was set to 120,000, and the MS2 resolution was set to 30,000. Precursor ions were fragmented by stepped high-energy collision dissociation (HCD) using stepped normalized collision energies of 20, 40, and 55%. Dynamic exclusion was enabled with an exclusion duration of 6 s using a mass tolerance of ±6 ppm. The automatic gain control (AGC) target value was set to 4×105 for the full MS and to 5×104 for the MS/MS spectral acquisition.

A blank (80% MeOH in H2O) sample as well as Mix1 were analysed throughout the sample set to check for mass error, retention time (RT) shift and overall signal reproducibility (defined as area under the curve, AUC). The results show high-quality data where the predefined acceptance criteria (< 5ppm for mass accuracy, 0.1 min for RT shift and <10% for AUC reproducibility) for these parameters were fulfilled.

MzMine was utilized to generate a table of features, which was successively manually curated to verify the correct feature ID from automatic open access and in-house library database match (in both MS1 and MS2).

### Proteomics analysis

Proteomics was performed at the Stein Family Mass Spectrometry Center in the Silberman Institute of Life Sciences, Hebrew University of Jerusalem.

### Pellet preparation

Single colonies were inoculated in 3 mL LB medium (with added kanamycin for mutants) and grown overnight at 37°C. Pre-cultures were washed three times with 1 mL 1xMSgg medium lacking CaCl₂ and FeCl₃, then resuspended in 3 mL of the same medium. The washed pre-cultures were used to inoculate the main cultures (4 mL complete 1xMSgg, no kanamycin) at an initial OD₆₀₀ of 0.01–0.02, which were incubated at 30°C, 200 rpm until OD₆₀₀ ≈ 0.8. Three biological replicates per strain were harvested by centrifugation (6000 × g, 10 min), washed twice with room-temperature 1X PBS, and resuspended in a lysis buffer (PBS + 4% SDS). The pellets in the lysis buffer were sonicated (5-20 second intervals for 10 minutes) until the buffer was clear and could be easily pipetted without clogging, and incubated for 5 minutes at 95°C. The lysate was centrifuged at max speed for 20 min to pellet cell debris, and the supernatant was collected into a clean tube and kept at-80°C.

### Sample preparation

Protein samples were digested and cleaned using S-Trap microcolumns (Protifi, LLC, Huntington, NY). Proteins in TRIS-HCl 25 mM pH 8 with 4% SDS were reduced by the addition of DTT to a concentration of 10 mM and incubated for 30 min at room temperature. Samples were then alkylated in 55 mM iodoacetamide and incubated for 30 min at room temperature in the dark. Phosphoric acid was added to a final concentration of 1.2%. Methanol-TRIS buffer (90% MeOH, 10% TRIS 0.5 M pH 7.1) was added to the samples at a ratio of 6:1 (buffer:sample) and loaded onto S-Trap columns by centrifugation at 1,000 ✕ g until all the sample was loaded onto the column (∼1 min). Columns were subsequently washed twice with 150 µL Methanol-TRIS buffer at 4,000 ✕ g. Sequencing-grade modified trypsin (Promega Corp., Madison, WI) was loaded onto the column (1 µg per column) and incubated at 47 °C for 90 minutes.

Peptides were eluted from the column, acidified, and desalted on homemade C18 stage tips ^71^. From the resulting peptides 0.35 µg (determined by Absorbance at 280 nm) from each sample was injected into the mass spectrometer for analysis.

### LC-MS/MS analysis

Proteomics analyses were performed using a Q Exactive-HF mass spectrometer (Thermo Fisher Scientific) coupled to an Ultimate 3000 Dionex (Thermo Fisher Scientific) UHPLC system. Peptides were separated on a 120 min acetonitrile gradient run at a flow rate of 0.30 mL min^-1^ on a reverse-phase 25-cm-long C18 column (Aurora Ultimate XT 25×75, ionopticks, AU). Survey scans (300–1,650 m/z, target value 3E6 charges, maximum ion injection time 20 ms) were acquired and followed by higher-energy collisional dissociation (HCD) based fragmentation (normalized collision energy 27). A resolution of 60,000 was used for survey scans, and up to 15 dynamically chosen most abundant precursor ions, with “peptide preferable” profiles were fragmented (isolation window 1.6 m/z). The MS/MS scans were acquired at a resolution of 15,000 (target value 1E5 charges, maximum ion injection times 25 ms). Dynamic exclusion was 20 sec. Data was acquired using Xcalibur software (Thermo Scientific). To avoid carryover, the column was washed with 80% acetonitrile, 0.1% formic acid for 25 min between samples.

### MS data analysis

Mass spectra data were processed using the MaxQuant computational platform, version 2.6.2.0. Peak lists were searched against two *B. subtilis* proteomes (UP000001570 and UP000306535) supplemented with custom sequences. The search included cysteine carbamidomethylation as a fixed modification, N-terminal acetylation and methionine oxidation as variable modifications, and allowed up to two miscleavages. The ‘match-between-runs’ option was used. The required FDR was set to 1% at the peptide and protein level. Relative protein quantification in MaxQuant was performed using the label-free quantification (LFQ) algorithm. MaxLFQ allows accurate proteome-wide label-free quantification by delayed normalization and maximal peptide ratio extraction_72._

### Differential proteomic analysis

Differential proteomic analysis was performed comparing *B. subtilis* NCIB3610 wild-type to Δ*ilvA*_ThrC(T55M), and Δ*cysE*_MetA(A28P/A108V) strains. Protein groups output tables were imported into Perseus (v2.1.3.0) for statistical analysis. Reverse hits, proteins identified only by site, and potential contaminants were removed prior to analysis. LFQ intensity values were log₂-transformed, and proteins with fewer than three valid values in at least one experimental group were excluded. Missing values were imputed from a normal distribution using the default Perseus settings (width = 0.3, down shift = 1.8). Differential protein abundance was assessed using a two-sided Student’s t-test with permutation-based FDR correction (FDR = 0.05, S₀ = 0.1) ^73^. The corresponding Perseus significance curve was used to classify proteins as significantly up-or downregulated in the volcano plot.

### Functional enrichment analysis

Significantly differentially abundant proteins identified by Perseus were analyzed for Gene Ontology (GO) term and KEGG pathway enrichment using ShinyGO ^74^, DAVID ^75^, and STRING-db ^76^ to compare enriched biological processes, molecular functions, cellular components, and metabolic pathways. The *Bacillus subtilis* 168 genome served as the background reference, and terms with an adjusted *P* value < 0.05 were considered significant.

### Calculation of ΔΔG values for mutations

To estimate the effect of mutations on protein stability, we used FoldX 5.1 ^77^. Input structures were chosen from experimentally solved PDB structures, or from experimentally solved structures of homologous proteins when the mutated position was resolved; when no experimental coordinates covered the site, a full-length AlphaFold model was used. Structures used were ThrC (PDB: 6NMX and 6CGQ ^78^), MetA (PDB: 2GHR ^79^), SerA (AlphaFold DB: AF-P35136-F1-v4), and CymR (AlphaFold DB: AF-O34527-F1-v4) ^80^. First, the experimental PDB structures were repaired to add missing atoms and optimize side chains using the RepairPDB command. Mutations were specified in FoldX individual list format (e.g., T55M;) in individual_list.txt. Mutations were then introduced with the BuildModel command using the default conditions. Per-run ΔΔG values were obtained from the Dif_* output files (total energy value). FoldX energies are reported in kcal·mol⁻¹ and follow the convention ΔΔG = ΔG(mutant) − ΔG(WT) (positive = destabilizing).

## Acknowledgements

The work by SH, SS, ABH, RM and LNG was funded by the Israeli Science Foundation grant No. 1168/24. Work by PCM, CCP, DR and LH was supported by the Novo Nordisk Foundation with Grant NNF20CC0035580. PCM, DR and LH are also supported by the Novo Nordisk Foundation with grant NNF24SA0100980.

## Notes

### Competing Interest Statement

The authors have declared no competing interest.

