## Supplementary Information for "Species-specific metabolic networks shape evolutionary routes to functional rescue"

[1] Department of Plant Pathology and Microbiology. Institute of Environmental Sciences. Faculty of Agriculture, Food and Environment. Hebrew University of Jerusalem, Israel

[2] Novo Nordisk Foundation Center for Biosustainability, Danmarks Tekniske Universitet, Kongens Lyngby, Denmark

* These authors contributed equally to this work

[+] Corresponding author

### List of tables

**Table S1:** Description of the 17 missing functions (below)

**Table S2:** Success & Growth Rate

**Table S3:** Similar EC numbers

**Table S4:** Mutation analysis

**Table S5:** The 40 acyl-transferases (E.C. 2.3.1.-) encoded by Bacillus subtilis

**Table S6:** UniProt Sequence ID

**Table S7:** Metabolomics

**Table S8:** Proteomics

**Table S9:** Strains and oligos used in this study

**File S1:** Coverage plots

**Table S1. The 17 *Bacillus subtilis* single-gene deletions identified as auxotrophic by Koo et al. (2017) used in this study, along with their corresponding missing enzyme function descriptions.**

Genes that were successfully bypassed in this study:

| | Function | E.C. number | Pathway and Reaction |
|---|---|---|---|
| *argH* | ArgH: Argininosuccinate lyase | 4.3.2.1 | Arginine biosynthesis;<br>L-argininosuccinate → arginine + fumarate |
| *aroC* | AroC: 3-dehydroquinate dehydratase | 4.2.1.10 | Aromatic amino-acids biosynthesis;<br>3-dehydroquinate → dehydroshikimate + $H_2O$ |
| *cysE* | CysE: Serine O-acetyltransferase | 2.3.1.30 | Cysteine biosynthesis;<br>serine + acetyl-CoA → O-acetyl-serine + coenzyme A |
| *hom* | Hom: Homoserine dehydrogenase | 1.1.1.3 | Methionine and threonine biosynthesis;<br>aspartate semialdehyde + NAD(P)H +$H^+$ → homoserine + $NAD(P)^+$ |
| *ilvA* | IlvA: Threonine dehydratase | 4.3.1.19 | Branched-chain amino-acids biosynthesis;<br>Threonine → 2-oxobutyrate + ammonium |
| *leuB* | LeuB: 3-isopropylmalate dehydrogenase | 1.1.1.85 | Leucine biosynthesis;<br>3-Isopropylmalate + $NAD^+$ →<br>2-keto-isocaproate + $CO_2$ + $NADH_2$ |
| *metA* | MetA: Homoserine O-succinyltransferase | 2.3.1.46 | Methionine biosynthesis;<br>homoserine + acetyl-CoA →<br>O-acetyl-homoserine + coenzyme A |
| *pheA* | PheA: prephenate dehydratase | 4.2.1.51 | Phenylalanine biosynthesis;<br>$H^+$ + prephenate → 3-phenylpyruvate + $CO_2$ + $H_2O$ |
| *serB* | SerB: Phosphoserine phosphatase | 3.1.3.3 | Serine biosynthesis;<br>phosphoserine + $H_2O$ → serine + phosphate |

Genes that were unsuccessfully bypassed in this study:

| | | | |
|---|---|---|---|
| *aroB* | AroB: 3-dehydroquinate synthase | 4.2.3.4 | Aromatic amino-acids biosynthesis;<br>3-deoxy-D-arabino-heptulosonate 7-phosphate → 3-dehydroquinate + phosphate |
| *hisC* | HisC: Histidinol-phosphate, | 2.6.1.9 | Histidine and aromatic amino-acids |

| | | | |
|---|---|---|---|
| | Tyrosine, and Phenylalanine aminotransferase | | biosynthesis;<br>3-phenylpyruvate + glutamate → phenylalanine + 2-oxoglutarate ;<br>4-hydroxyphenylpyruvate + glutamate → tyrosine + oxaloacetate |
| *ilvD* | IlvD: Dihydroxy-acid dehydratase | 4.2.1.9 | Branched-chain amino-acids biosynthesis;<br>2,3-dihydroxyisovalerate → 2-ketoisovalerate + $H_2O$ ;<br>2,3-dihydroxy-3-methylvalerate → 2-oxo-3-methylvalerate + $H_2O$ |
| *metE* | MetE: Methionine synthase | 2.1.1.14 | Methionine biosynthesis;<br>L-homocysteine + 5-methyltetrahydropteroyl-triglutamate → methionine + tetrahydropteroyl-triglutamate |
| *pyrF* | PyrF: Orotidine 5-phosphate decarboxylase | 4.1.1.23 | Pyrimidine biosynthesis;<br>orotidine-5'-P + $H^+$ → UMP + $CO_2$ |
| *serA* | SerA: Phosphoglycerate dehydrogenase | 1.1.1.95 | Serine biosynthesis;<br>3-phosphoglycerate + $NAD^+$ → phosphohydroxypyruvate + $H^+$ + $NADH_2$ |
| *sirB* | SirB: Sirohydrochlorin ferrochelatase | 4.99.1.4 | Siroheme biosynthesis, sulfite reduction;<br>$Fe^{2+}$ + sirohydrochlorin → 2 $H^+$ + siroheme |
| *thrC* | ThrC: Threonine synthase; minor threonine dehydratase (IlvA) activity | 4.2.3.1 | Threonine biosynthesis;<br>Threonine → 2-oxobutyrate + ammonium ;<br>O-phosphohomoserine + $H_2O$ → threonine + phosphate |

### Supplementary Figures

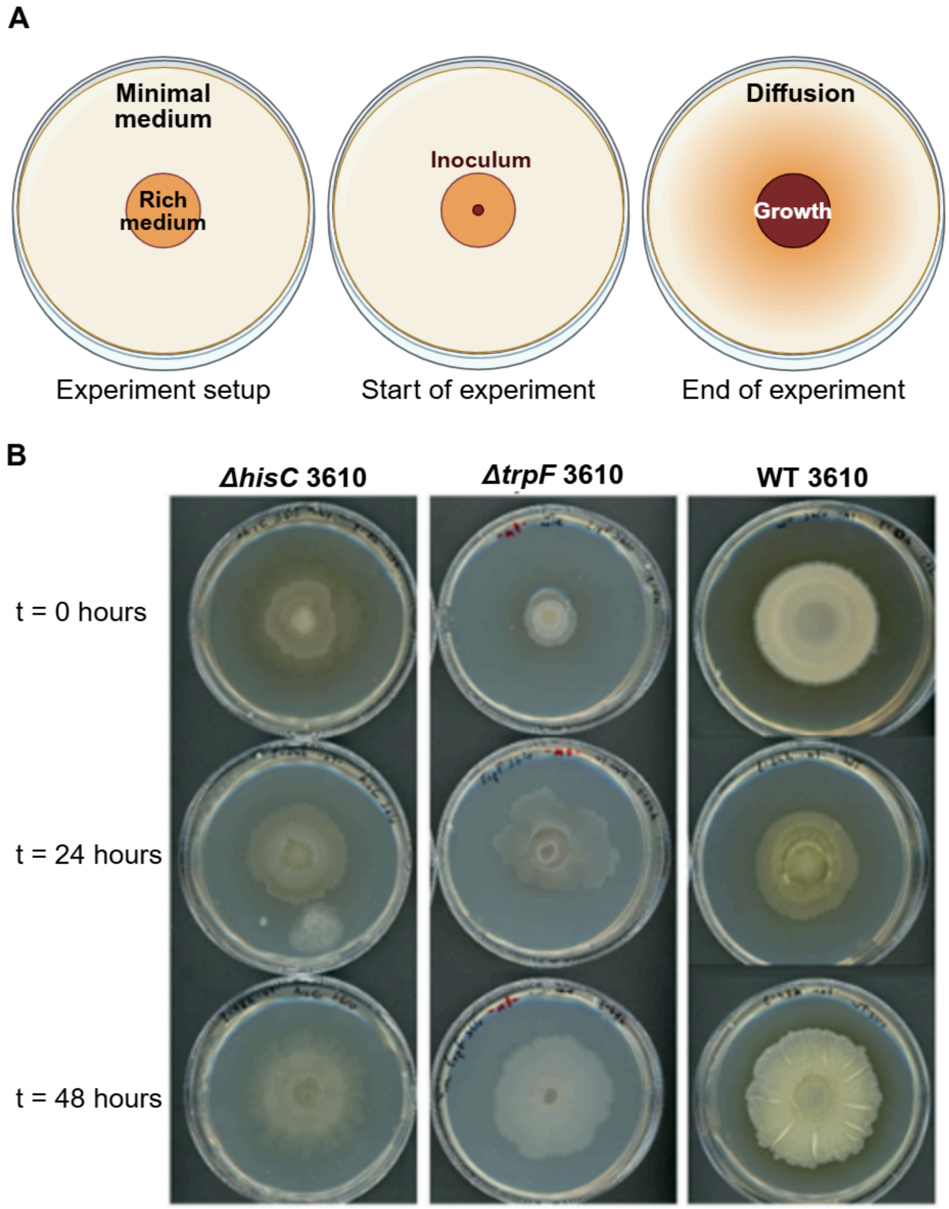


**Figure S1. Gradient plate design and calibration. (A)** A schematic representation of the gradient environments and their preparation. An MSgg agar plate is prepared. Once solidified, a cavity is created in the center and filled with LB agar. Once the entire plate had dried, we inoculated its center with 5 μl of a pre-cultured auxotrophic strain. While the cells grow, the LB nutrients diffuse toward the MSGG, creating a nutrient gradient. (**B)** Examples from the triple time point gradient evaluation. The strain genotype is stated at the top of each triplet; incubation time before inoculation is on the left. All plates were prepared on the same day, but inoculated within a 1-day window of one another. Pictures were also taken on the same day.

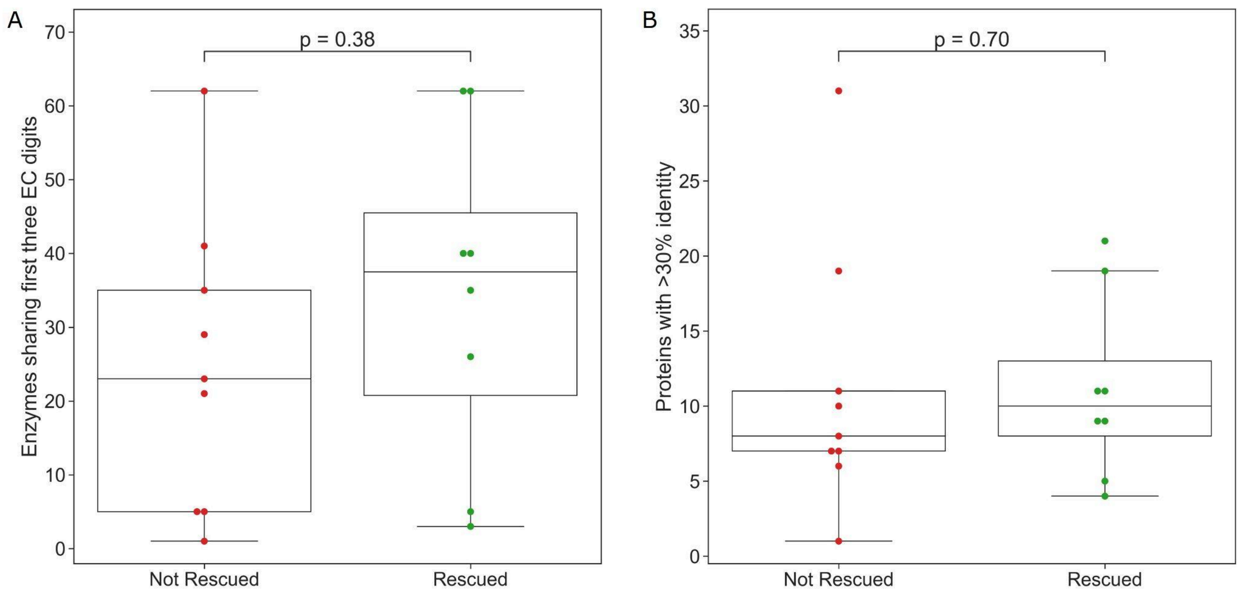


**Figure S2. The number of enzymes similar in function or sequence to the knocked-out gene does not predict metabolic bypass.** Distributions of the number of **(A)** Enzymes in the same EC sub-subclass (sharing the first three digits) and **(B)** proteins in the genome sharing at least 30% identity for the knocked-out gene of each auxotroph. We split the data into auxotrophs that were rescued at least once and those that were not rescued from extinction. Dots represent individual auxotrophs, solid lines represent the median, boxes represent the interquartile range, and whiskers are expanded to include values no further than 1.5× the interquartile range. Mann–Whitney–Wilcoxon two-sided tests were performed, and p-values are shown on each graph.

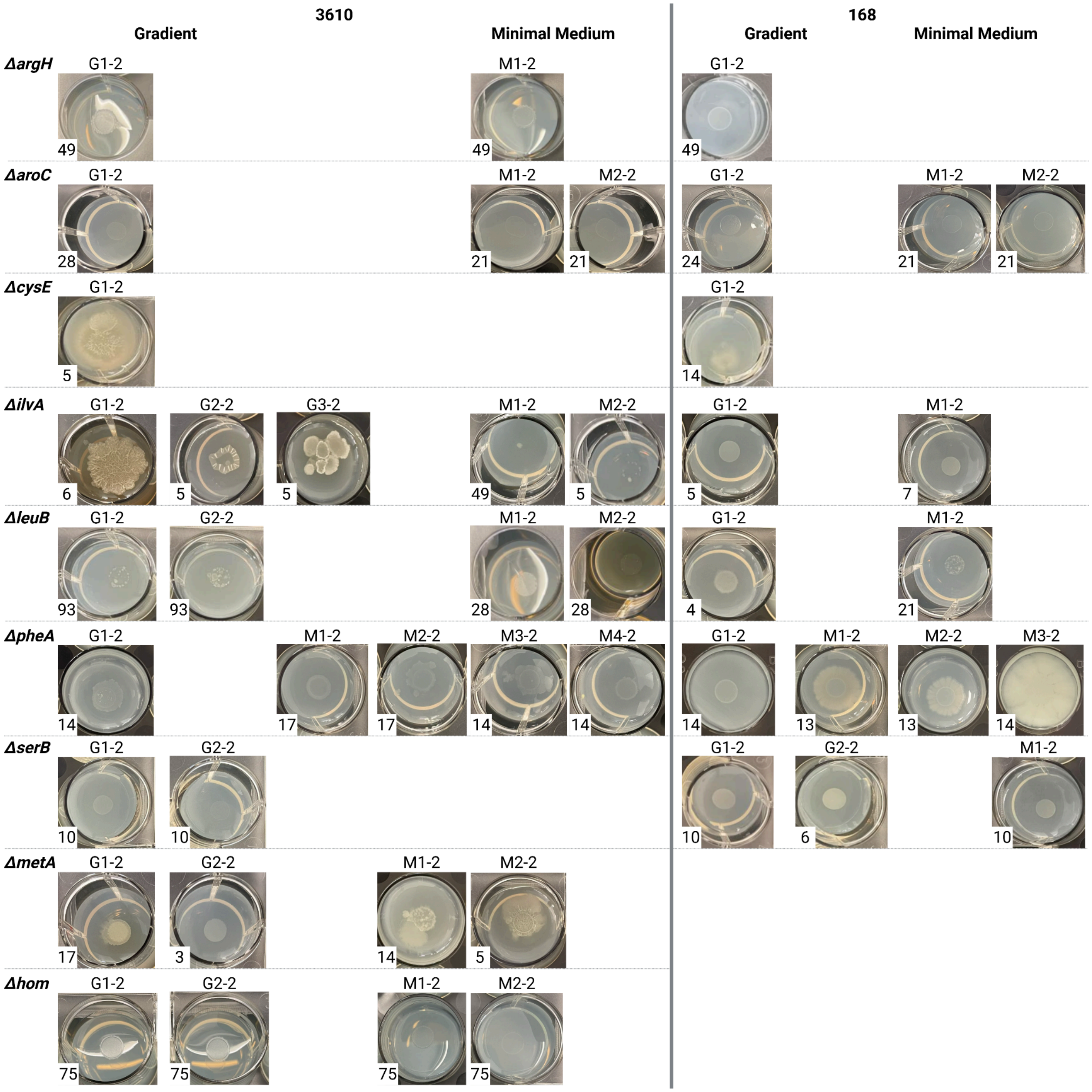


**Figure S3. The 46 evolved populations after their second round of growth in MSgg.** Pictures were taken after the indicated number of days (bottom left). The population name is indicated. G means the experiment started with a gradient plate, while M indicates the experiment started in minimal medium. Numbers were used when more than one population evolved under the same condition. The number after the hyphen indicates the MSgg cycle. Here, the number 2 is shown for all populations, indicating the second passage of the ALE process.

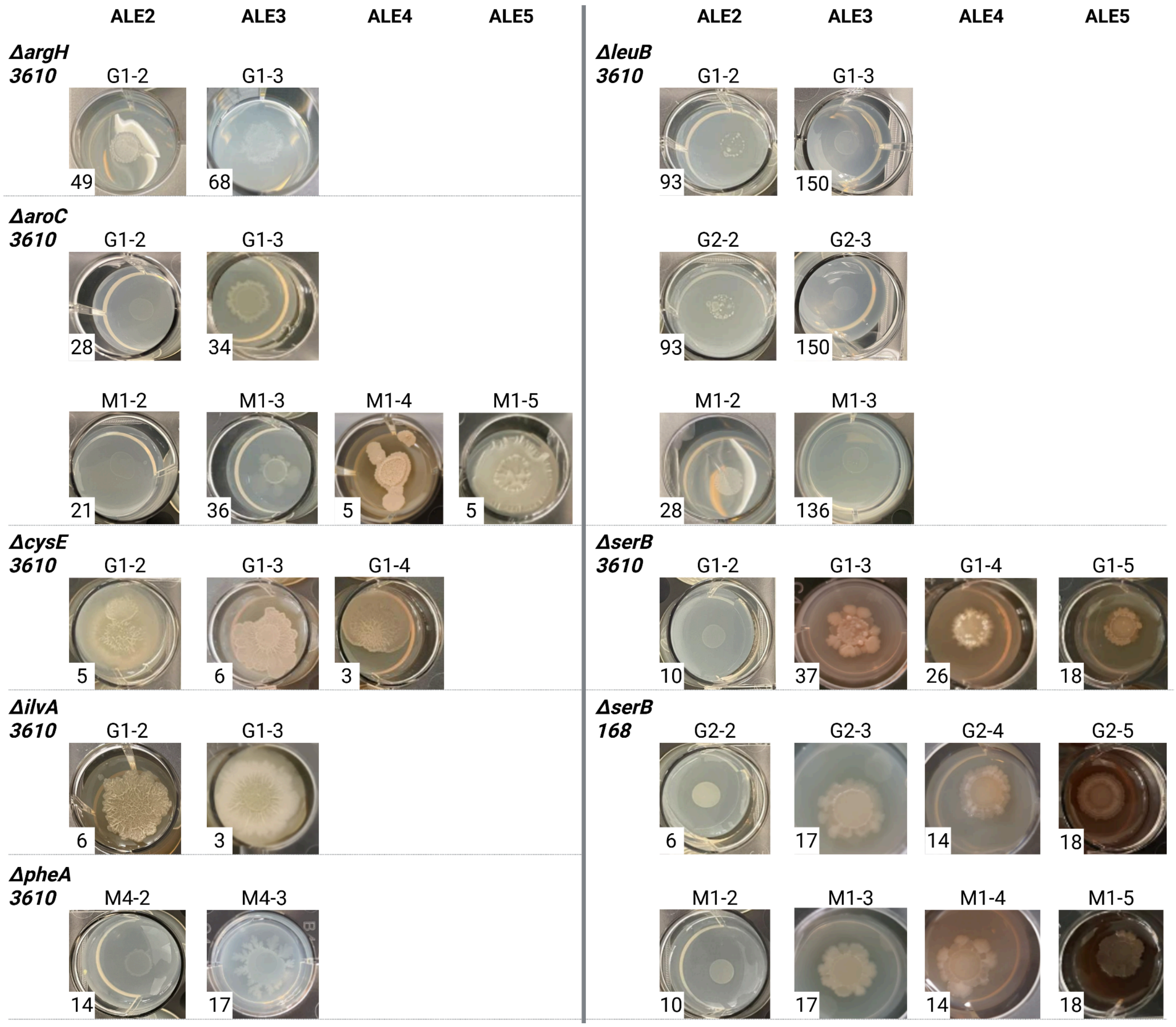


**Figure S4. Additional ALE rounds of 12 evolved populations.** The pictures were taken after the indicated number of days (bottom left). Population names are indicated. G means the experiment started with a gradient plate, while M indicates the experiment started in minimal medium. Numbers were used when more than one population evolved under the same condition. The number after the hyphen indicates the MSgg cycle.

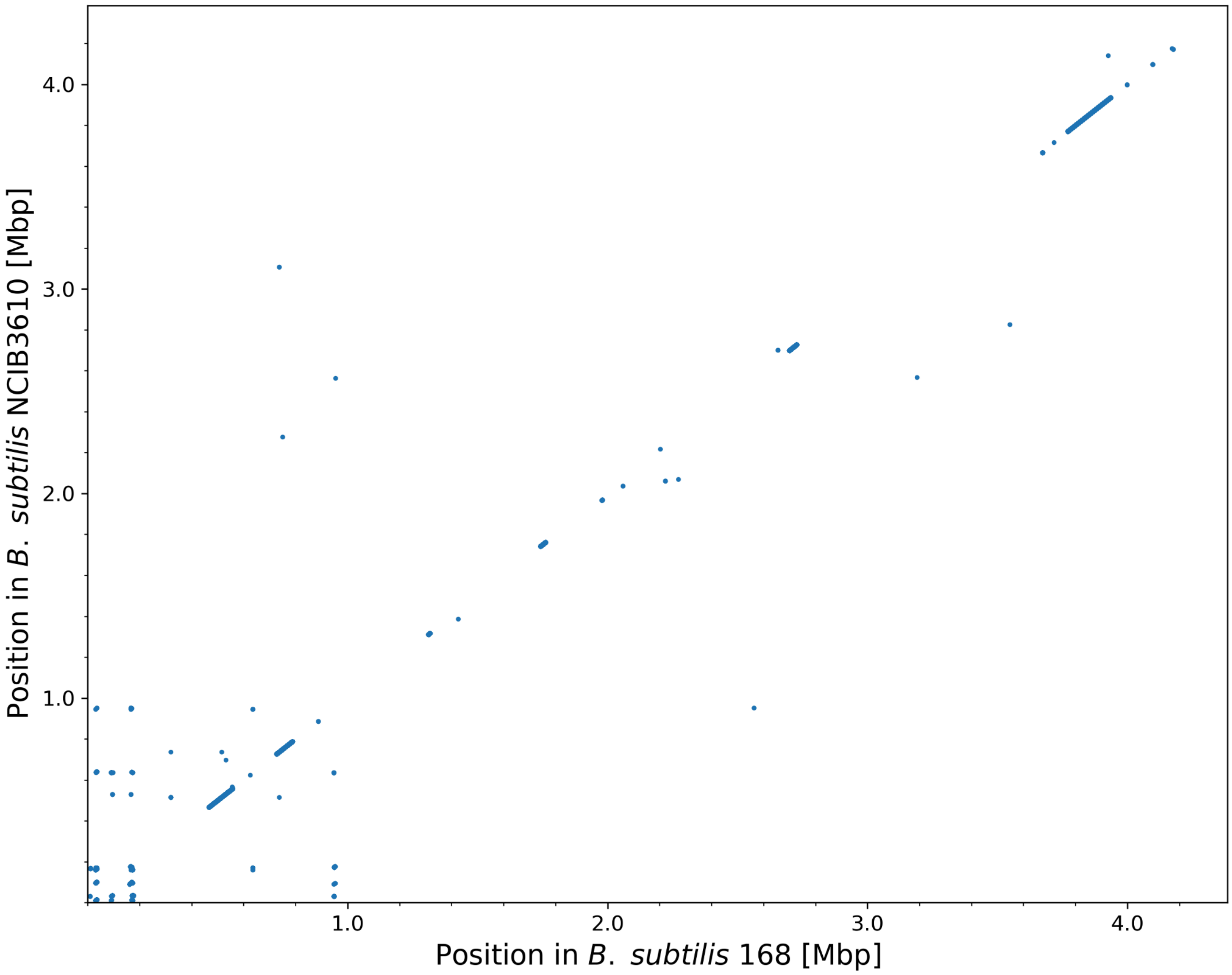


**Figure S5. Genome sequence similarity between *B. subtilis* strains 168 and NCIB3610.** Dot plot showing shared k-mer matches between *B. subtilis* strain 168 (x-axis) and NCIB3610 (y-axis). Each point represents a perfect match of a 20 bp k-mer (k=20) identified approximately every 10 bp along the *B. subtilis* 168 genome and mapped to the *B. subtilis* NCIB3610 genome. Both axes are scaled in million base pairs (Mbp). Diagonal alignment of dots indicates strong genome similarity (Pearson correlation coefficient r=0.998), i.e., the 20 k-mer fragments match in sequence and position. Deviations from that line reflect matching sequences that are found at different positions along the two genomes.

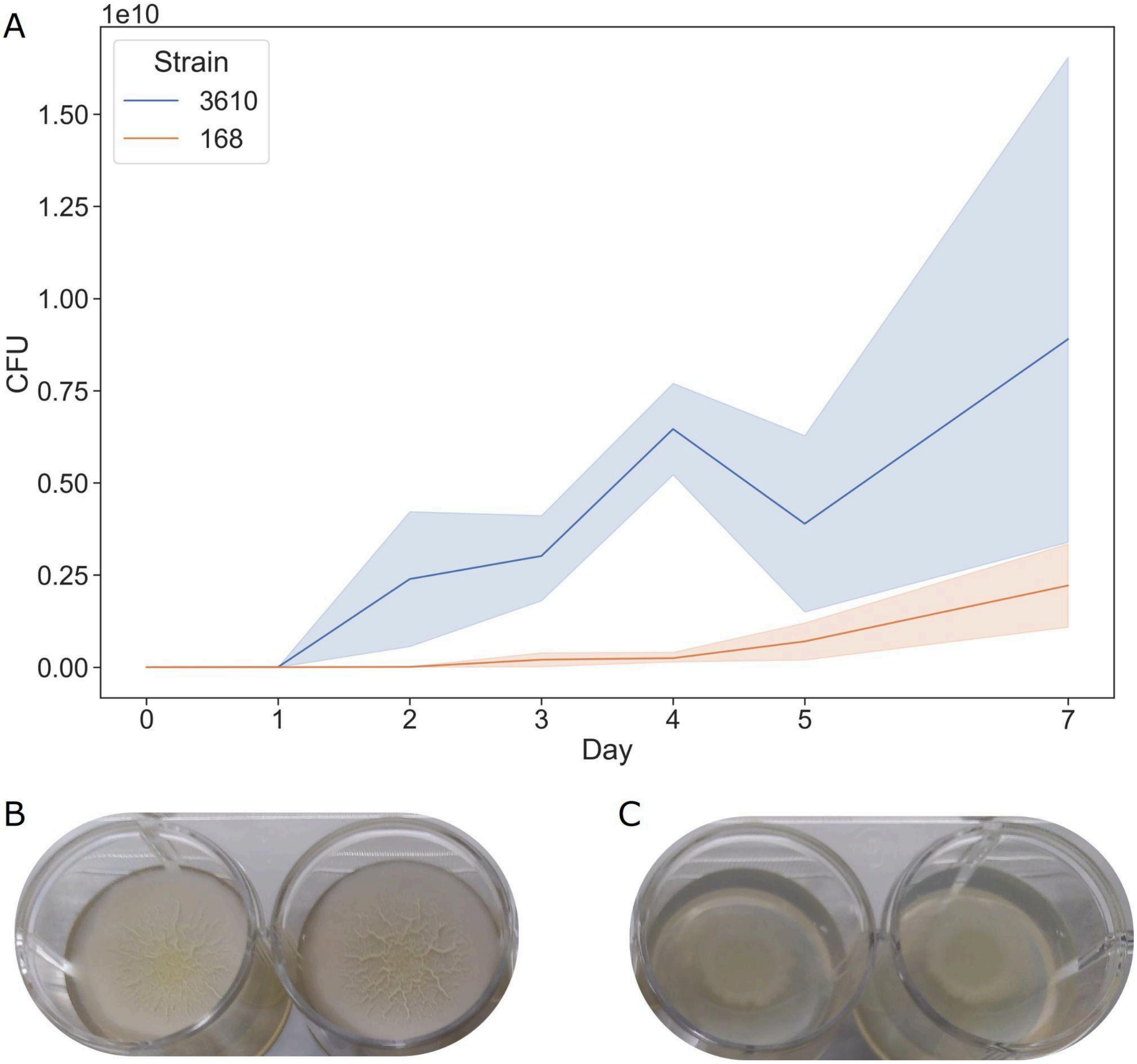


**Figure S6. Growth of *B. subtilis* 168 and 3610 in the MSgg medium. (A)** Comparison of the growth rate of *B. subtilis* NCIB3610 and 168. Colony-forming units (CFU) over 7 days for strains 3610 (blue) and 168 (orange). The solid line represents the mean, and the shaded area the 95% confidence interval. Exponential-phase data (days 0–2 for 3610; 0–3 for 168) were fitted by linear regression to estimate doubling times: 3610=1.83 ± 0.19 h and 168=3.35 ± 0.65 h, indicating that 3610 grows 1.83 times as fast. **(B–C)** Representative day-7 Biofilms. Despite having similar final CFU counts (Mann–Whitney–Wilcoxon two-sided test, $P$ = 0.11), 3610 populations are visibly larger and more spread out than those of 168.

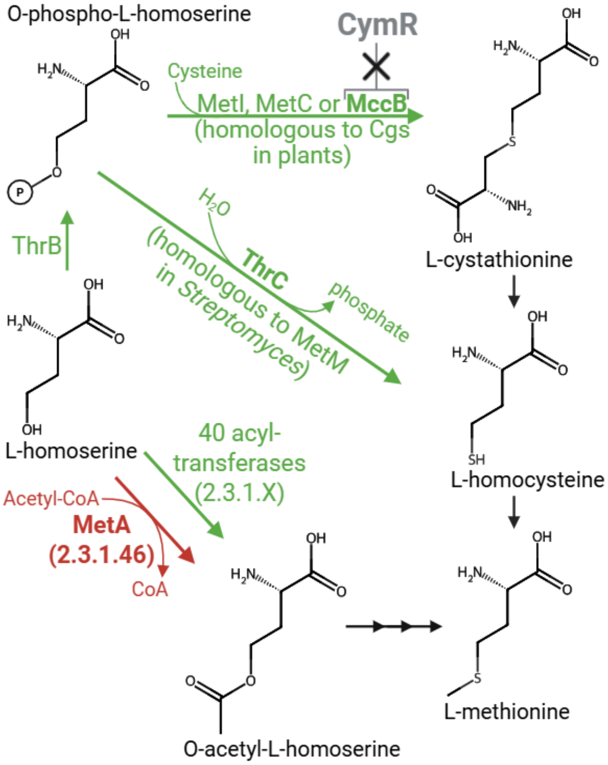


**Figure S7. A simplified schematic representation of the suggested methionine biosynthetic pathways bypassing the function of *metA*.** Methionine biosynthesis from L-homoserine through its conventional route (red) and three proposed bypasses (green), with their route number mentioned in a circle. Enzymes are named by their 4-letter code: MccB, cystathionine lyase/ homocysteine gamma-lyase; MetA, homoserine O-succinyltransferase; MetC, cystathionine beta-lyase; MetI, cystathionine gamma-synthase/O-acetylhomoserine thiolyase; ThrB, homoserine kinase; ThrC, threonine synthase; MetM, homocysteine synthase; Cgs, cystathionine γ-synthase.

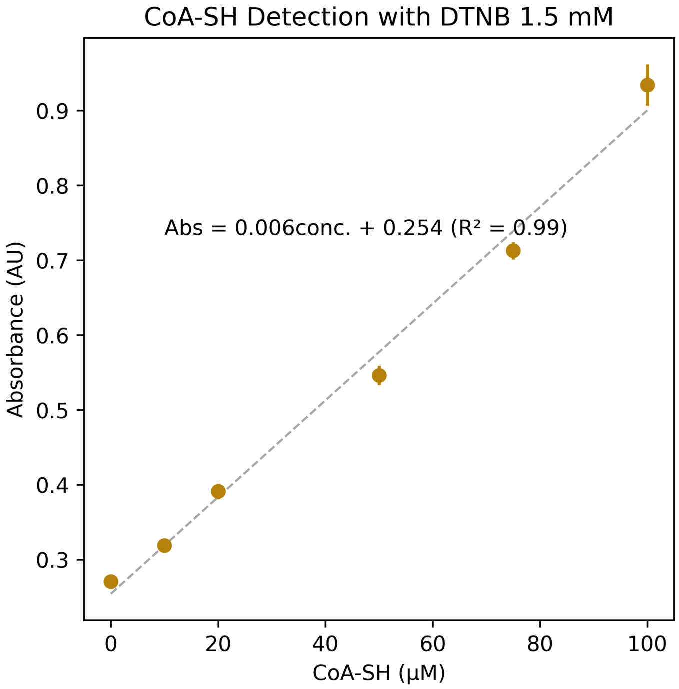


**Figure S8. CoA-SH standard calibration curve for the MetA activity assay (related to Figure 4C).** Absorbance was measured using increasing concentrations of CoA-SH in the presence of 1.5 mM DTNB. Points represent measured absorbance values, and the dashed line represents the linear regression. The regression equation and coefficient of determination ($R^2$) are shown.

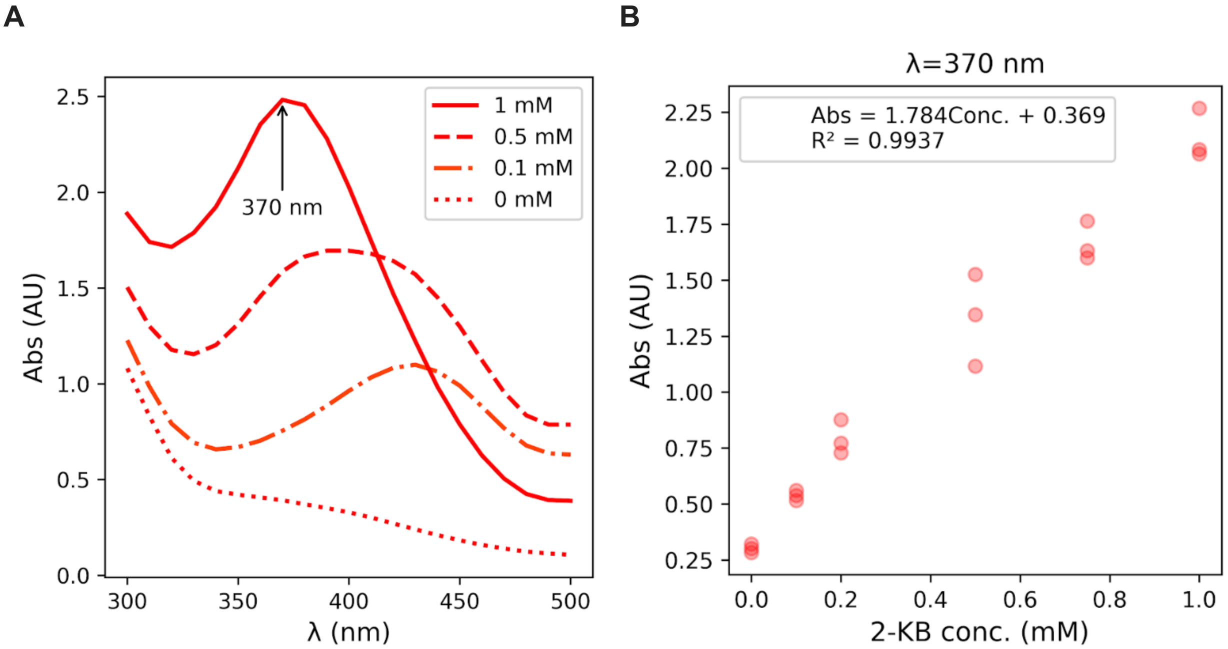


**Figure S9. 2-Oxobutanoate detection and calibration for the ThrC activity assay (related to Figure 5B). (A)** Absorbance spectra of 2-oxobutanoate standards at the indicated concentrations. The selected detection wavelength (370 nm) is indicated. **(B)** Standard calibration curve relating 2-oxobutanoate concentration to absorbance at 370 nm. The linear regression equation and coefficient of determination ($R^2$) are shown.
